# Coupling cell growth to protein secretion enables pooled screening at single-cell resolution in bioreactors

**DOI:** 10.64898/2026.09.14.751527

**Authors:** Henri Galez, Alicia Franja Da Silva, Cecilia Capela, Sara Napolitano, Gregory Batt

**Affiliations:** Institut Pasteur, Inria, Université Paris Cité, Paris, France; IFP Energies nouvelles, Rueil-Malmaison, France; AgroParisTech, Palaiseau, France

## Abstract

Preserving the genotype-phenotype linkage is critical in high-throughput screens exploring large parameter spaces. Screening pooled libraries for recombinant protein secretion is therefore particularly difficult, since protein diffusion into the medium disrupts this linkage. Existing methods impose significant restrictions on the environment of the screened cells. Here, we present AutoGrowth, a growth-based screening strategy linking growth rate to secretion via a single-cell biosensor. We demonstrated the proportional modulation of growth rate by secretion and validated the approach on five model proteins. As proof of concept, we screened a signal peptide library in bioreactors for improved secretion of a single-chain variable fragment by coupling AutoGrowth with deep sequencing. Additionally, through spike-in experiments, we showed that AutoGrowth can identify favorable variants diluted at ratios as low as one in hundreds of millions in three days. Our growth-based strategy provides unprecedented throughput potential and ensures selection of genetic variants presenting advantageous secretion and fitness capabilities.

## Introduction

Recombinant proteins are of major importance in the pharmaceutical and industrial sectors^1,2^, and their role is expected to expand further into the food industry through precision fermentation^3^. These proteins are produced in mammalian or microbial cells, the most common microbial cell factories being *E. coli*, *B. subtilis*, *P. pastoris*, and *S. cerevisiae*^4^. The latter three offer the advantage of secreting the recombinant protein of interest (POI) into the culture medium, thereby facilitating downstream operations. Among these, *S. cerevisiae* is an attractive platform combining industrial robustness with extensive genetic tractability.

Optimizing recombinant protein secretion requires engineering both the chassis strain and the expression system. The former is typically engineered by introducing modifications of the endogenous genetic network, while the latter is typically optimized by finding the best combination of several parameters, among which promoter, signal peptide (SP), codon usage, and gene dosage^5^. This represents a very large parameter space. Moreover, combinations improving secretion are to a large extent POI-specific^6,7^. Efficiently navigating this parameter space can be achieved through the screening of pooled libraries, an approach greatly facilitated by molecular biology tools, such as Modular Cloning and CRISPRi/a. They greatly facilitate building large pooled libraries by enabling one-pot combinatorial assemblies^8^ and genome-wide perturbations^9^.

Such large-scale libraries require high throughput screening methods, scaling beyond traditional pipelines based on 96-well plate format^10–15^. Methods assessing the secretion capacity of individual cells within a cell culture have much higher throughput. Single-cell encapsulation into micro-droplets allows such individual evaluation and offers high throughput sorting capacity, up to 2×10^6^ droplets per screen, through fluorescence-associated droplet sorting (FADS)^9,16–22^. However, these assays are mostly adapted to proteins having specific enzymatic activities. Moreover, cells are screened in a micro-environment different from what they would encounter in fermentation processes^23,24^. Other methods exploit limited protein diffusion on solid medium to identify colonies with a secretion-indicative phenotype^25–27^. They suffer from the same limitations of enzymatic activity-dependent readouts and screening conditions incompatible with bioproduction contexts^28^. Methods based on secrete-and-capture^29,30^ also require particular conditions, static culture or high viscosity, to prevent protein diffusion. Culture conditions influence cell physiology and metabolism, which in turn affect secretion capacity^31^. Screening libraries under conditions closer to fermentation could yield results more likely to translate to large-scale processes. Screening pooled libraries in fermentation-like liquid culture brings two challenges: quantifying the secretion capacity of each cell despite protein diffusion (maintenance of the genotype-phenotype linkage) and selecting for high secretors. Methods based on yeast surface display (YSD) address these challenges by retaining the POI on the cell wall through fusion with an anchor and selecting variants with enhanced display through fluorescence-associated cell sorting (FACS)^32,33^. This approach rests on the assumption that secretion capacity of a cell correlates with the amount of POI retained on its cell wall. This is not always true. For example, the Aga2 anchor has been shown to interfere with secretion for several POIs, precluding screening for improved secretion of these POIs^33^. In addition, methods based on FACS screen up to ∼10^7^ cells, which limits the library size to ∼10^5^ variants, assuming a 100x coverage. While this is sufficient in many cases, the size of modern libraries can reach up to 10^8^ variants^34^. Collectively, these technologies have greatly advanced our capacity to optimize POI secretion in yeast. Yet, they fall short of supporting large POI-agnostic screens for secreted proteins under conditions approaching those of fermentation.

Here, we present the so-called AutoGrowth strategy: a single-cell biosensor that couples growth to recombinant protein secretion. We demonstrate that our biosensor possesses three key properties: it allows a cell to sense its own secretion despite protein diffusion; it shows monotonic and stable activation for various POIs; it enables the modulation of the cell growth rate. As proof of concept, we screened in bioreactors a small library of signal peptides for increased secretion of a single-chain variable fragment. Coupling AutoGrowth-based enrichments with deep sequencing allowed quantitative assessment of the performance of each signal peptide. For applications to larger libraries, we demonstrated with spike-in experiments the capacity to identify scarce good variants, one in a hundred million, in only three days of selection. This is the only method capable of screening very large libraries for improved secretion of a protein of interest in standard culture conditions. Critically, it requires no specific enzymatic activity, and further, accounts for the fitness of the selected variants.

## Results

### AutoGrowth strategy and genetic designs for a growth-based secretion biosensor

Improving secretion of a recombinant POI in yeast is done by tuning many parameters, resulting in large parameter spaces that can be explored through the screening of pooled libraries (Figure 1A). Such screenings come with the challenge of measuring secretion capacity at the single-cell level and being able to select for it. We developed a single-cell biosensor for secretion through an autocrine signaling, in which secreted proteins trigger downstream activation conferring a growth advantage. This biosensor enables enrichment of good genetic variants by simply propagating the pooled library.

**Figure 1:**
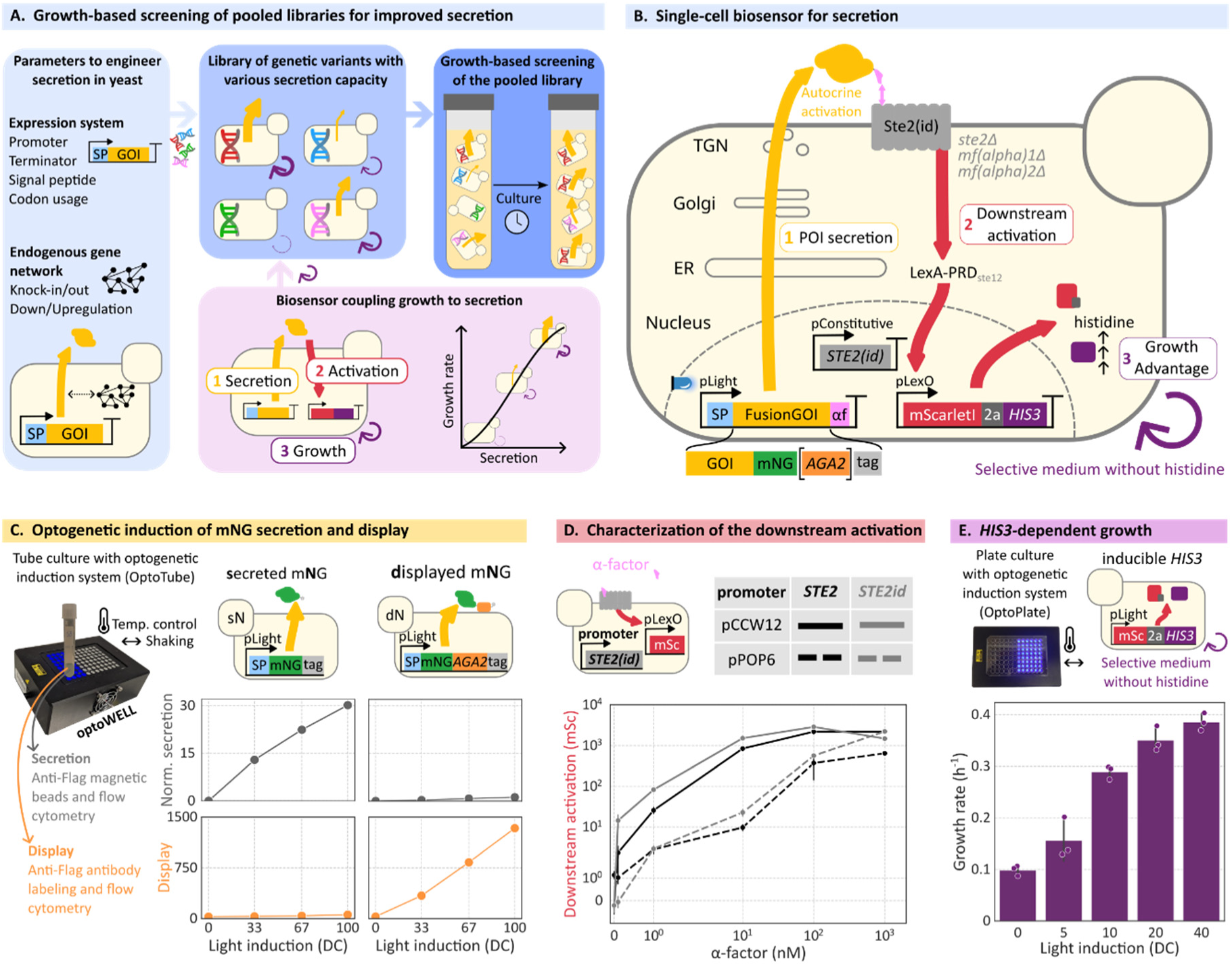
AutoGrowth strategy to couple growth to secretion at the single-cell level. A) Library screening with the AutoGrowth strategy. Efficient exploration of the parameter space is achieved by building libraries of genetic variants. Building them in a chassis containing the biosensor coupling growth to secretion enables growth-based screening. **B)** Molecular description of the biosensor. The POI is fused at its C-terminal with the 13-residue α-factor, unique requirement on the POI cassette, and in our case, expressed from a light inducible promoter for precise, optogenetic induction. For convenience, to track and to quantify the selected POI, we fuse it to a tagged mNeonGreen (mNG). We also studied the POI display by fusing it with the Aga2 anchor. The Ste2 receptor and the synthetic transcription factor LexA-PRD_Ste12_ are expressed from constitutive promoters. The synthetic promoter LexO6x pLeu2m drives the expression of the mScarletI.2a.*HIS3* reporter. Hijacking of the Ste2/α-factor pathway is made possible thanks to the engineered background strain yWS677^35^. **C)** Assessment of the POI module. Strains secreting or displaying mNG are grown under various duty cycles of blue light illumination using the OptoWELL device. After 10 hours of culture, secretion and display are measured for both strains. Secretion is measured using anti-Flag magnetic beads capturing secreted mNG, whose fluorescent signal is then measured by flow cytometry of the beads. Display is measured by labeling cells with a primary anti-Flag bound by a secondary antibody conjugated to a blue fluorescent dye, followed by flow cytometry of the cells. For secretion and display, we represent the median fluorescence of the distribution of beads and cells, respectively. **D)** Assessment of the sensing module. Cultures of strains expressing *STE2* or *STE2id* from the pPOP6 or pCCW12 promoters are supplemented with various concentrations of α-factor, and cell fluorescence is assessed after 10 hours by flow cytometry. Experiments are done in biological triplicate; we represent the mean and the standard deviation. **E)** Growth module assessment. A strain expressing mScarletI.2a.*HIS3* from the pLight promoter is grown under varying light duty cycles for 12 hours in complete medium before inoculation of histidine-lacking medium. OD measurements are then taken at multiple timepoints for growth rate computation. Experiments are done in biological triplicate; we represent the mean and the standard deviation.

Concretely, we coupled POI secretion to growth by fusing the α-factor peptide to the POI and placing the auxotrophic marker *HIS3* downstream of a streamlined α-factor signaling pathway (Figure 1B). POI secretion allows α-factor to bind to its cognate receptor Ste2, which leads to downstream activation of an mScarletI.2a.*HIS3* reporter conferring growth advantage in selective medium. An engineered genetic background (*i.e.* chassis) allows us to hijack the yeast mating pathway in a controlled and orthogonal manner^35^. POI secretion, downstream activation, and *HIS3*-dependent growth were characterized quantitatively to gain confidence in our ability to combine them into the desired biosensor.

First, we tested whether our optogenetic induction system permits secretion and display of mNeonGreen (mNG) in the chassis (Figure 1C). mNG is used here as a model protein for an easy-to-secrete POI (see Supplementary Note 1 for the list of model proteins, plasmids, and strains used in this study). We used OD normalized secretion as a proxy for single-cell secretion capacity, to be compared later with downstream activation. Cells secreting mNG showed increasing secretion with induction, while cells displaying mNG had minimal secretion, indicating the efficient cell wall anchoring through Aga2. Consistently, cells displaying mNG exhibited a strong induction-dependent display signal, whereas cells secreting mNG showed none. Display signal was confirmed by confocal microscopy (Supplementary Note 2). In addition, the induction range did not saturate the secretion and display capacities of the chassis for the easy-to-secrete mNG. Yet, we noticed that full induction of secreted mNG slightly altered cell fitness (Supplementary Note 2).

Second, we assessed downstream activation with sensing modules containing different promoter-receptor pairs and different variants of Ste2 (Figure 1D). Tuning receptor expression allows adjustment of both sensitivity and dynamic range of the response. Activation of Ste2 leads to its internalization and desensitization^36^, notably in the case of autocrine activation^37^. Such regulation could affect the stability of the response negatively. We therefore included an internalization-deficient variant of the receptor, Ste2id, shown previously to be more present at the plasma membrane and more sensitive^38^. Both receptors exhibited similar response profiles, with Ste2id achieving moderately higher activation. Regarding promoters, while the strong promoter pCCW12 conferred higher sensitivity, while the weak promoter pPOP6 generated broader operational ranges for both receptors. Notably, the pPOP6-Ste2id module spanned both operational and dynamic ranges over four orders of magnitude.

Third, we tested the possibility of modulating the cell growth rate through the controlled expression of *HIS3* in the absence of histidine in the medium (Figure 1E). Here, *HIS3* expression is modulated by placing the mScarletI.2a.*HIS3* reporter directly under the light inducible promoter. Cultures with various optogenetic inductions (light duty cycles, DC) have been performed. Results showed that the various levels of *HIS3* expression caused graded growth rates, with 40% DC enabling cells to recover the usual yeast growth rate, above 0.35 h^−1^, in our conditions.

Successful secretion and display of mNG, a large sensing operational range, and tunable growth rates collectively support the feasibility of linking growth rate to POI secretion/display through an autocrine loop.

### Core autocrine device for quantifying self-secretion capacity

To screen pooled libraries, the biosensor must be “single-cell”, meaning that proteins secreted by a cell should only generate downstream activation of that cell (*i.e.* autocrine-only activation). A second necessary property is that downstream activation scales monotonically with secretion capacity, ensuring that differences in secretion are reflected in the response. Lastly, downstream activation must be stable over time to support growth-based screening over several days.

First, we investigated whether our device enables autocrine-only activation by assessing co-cultures of two strains (Figure 2A). The first one, named autocrine strain, contains both the POI-expressing module and the sensing module. The second strain, named paracrine strain for simplicity, contains only the sensing module. It constitutively expresses a blue fluorescent protein for strain differentiation and determination of strain-specific downstream activation. Four autocrine strains were tested, each pairing a POI expression system (secreted or displayed) with a sensing module configuration (high or low Ste2id expression). They were each co-cultured with the paracrine strain harboring the corresponding sensing module, yielding four co-cultures. Downstream activation was measured for both strains in each co-culture. For low sensitivity strains, displayed mNG (dNL) caused higher autocrine activation than secreted mNG (sNL). This was expected, since the Aga2 tag retains POI at the cell surface. For both high sensitivity strains, sNH and dNH, the activation levels are high enough to saturate the sensor response (Figure 1D). Therefore, we selected the low sensitivity strains for all subsequent experiments. No paracrine activation was observed in any of the co-cultures. The total absence of paracrine activation was not expected for sNL and sNH as secreted mNG released in the medium should be able to activate other cells. This could have come either from a local gradient of POI concentrations, higher around the autocrine cells, or from an internal activation of Ste2id, which also takes the secretory pathway to reach the cell surface. Using dTA, an α-factor antagonist, and a Golgi-cleavable protection, we showed that the activation of the Ste2id receptor takes place mostly at the cell surface (Supplementary Note 3). In addition, we found that an N-terminal fusion of α-factor to a simple Flag-HIS tag is sufficient to almost abolish paracrine activation (Supplementary Note 3), in agreement with the role of α-factor N-terminal moiety to modulate paracrine activation^39^. Thus, our biosensor design, and particularly the α-factor fusion to the POI, robustly supports autocrine-only activation, with the readout reflecting the cell surface localization of the POI.

**Figure 2:**
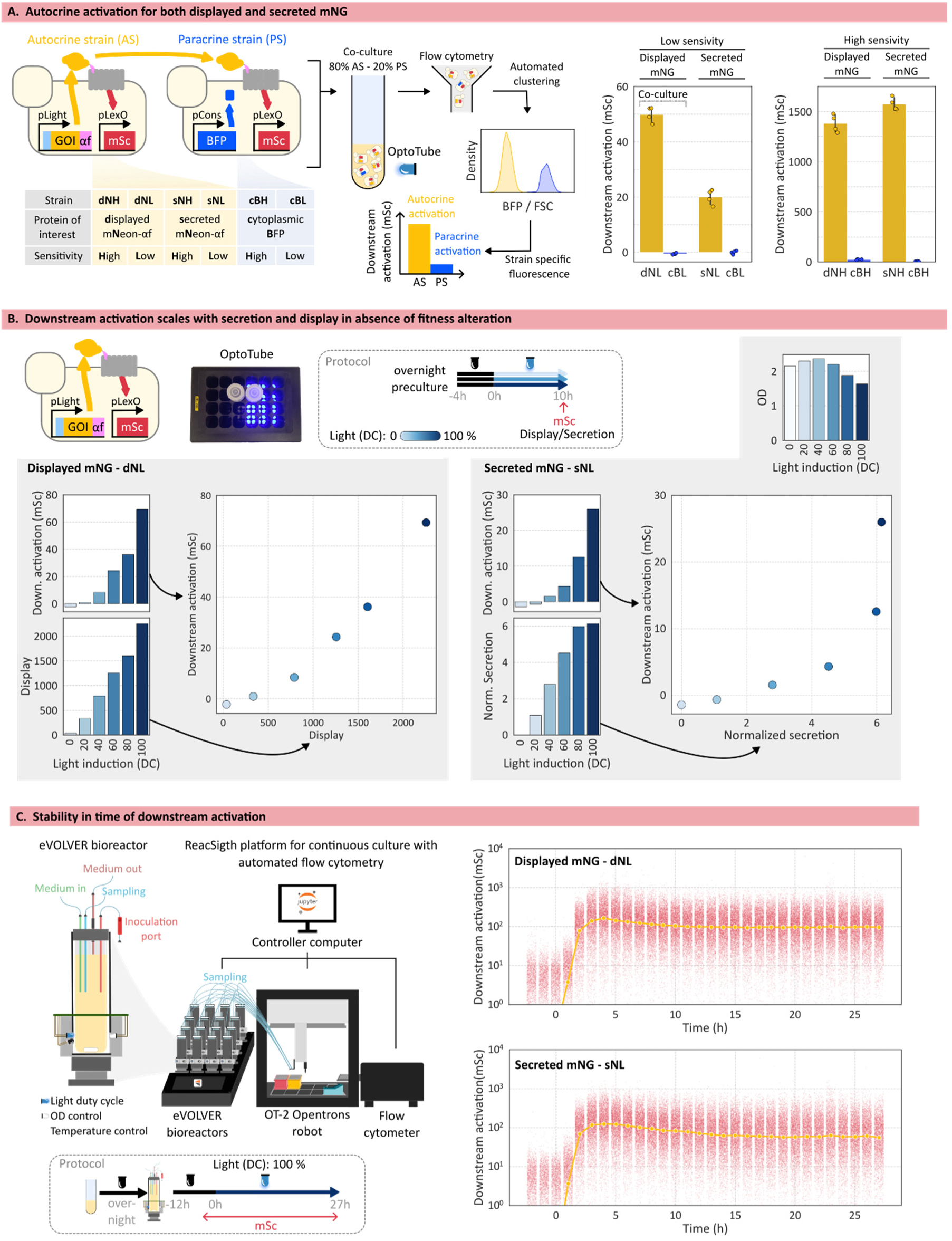
Downstream activation as a stable readout of single-cell secretion or display. **A)** The biosensor enables autocrine-only activation. Autocrine strains containing both a POI module, secreted/displayed mNG fused to α-factor, and a Ste2id-based sensing modules are co-cultured with an accessory strain containing only the sensing module. This second strain, named paracrine strain for simplicity, expresses cytoplasmic BFP for differentiation of the two strains using flow cytometry with automated clustering. Ste2id is expressed by pCCW12 or pPOP6 promoters, denoted “High” and “Low” sensitivity respectively. After 10 hours of culture under blue light, fluorescence of cells is measured by flow cytometry. Experiments are done in biological quadruplicate; medians of each population are represented by dots, and their mean and the corresponding standard deviation is shown with the bar plot. **B)** Downstream activation as a readout of secretion/display of mNG. Autocrine strains, secreting or displaying mNG, are grown under various light duty cycles for 10 hours before flow cytometry measurements of cell fluorescence, secretion (anti-Flag magnetic beads) and display (anti-Flag antibody labeling). Medians of distributions are represented. **C)** Assessment of biosensor activation stability. eVOLVER bioreactors are enhanced with automated real-time cytometry using the ReacSight strategy. Bioreactors are inoculated with autocrine strains, secreting or displaying mNG, and conducted in turbidostat regime (dilutions to maintain OD at 0.5). Cells are first grown in the dark to allow them to reach exponential growth before starting blue light illumination. Cell fluorescence is measured every hour with automated flow cytometry. Single-cell cytometry data are represented in red (with random artificial jitter on the x-axis), and the median in yellow.

Next, we sought to determine whether downstream activation is a pertinent proxy for secretion. We established the relationship between the two by applying a range of light DCs to secrete or display mNG at different levels. For dNL, downstream activation scaled with display (Figure 2B, left). For sNL, a saturation of secretion was observed above 80% DC (Figure 2B, right), consistent with previous results obtained using this optogenetic induction system^40^. Although mNG is efficiently secreted, strong induction seems to alter cell fitness, as shown by a lower final OD. Downstream autocrine activation scales with normalized secretion until the saturation threshold of 80%. In conclusion, downstream activation matches secretion or display in absence of altered fitness, suggesting a good applicability of the device for pooled library screening for improved secretion.

Lastly, the stability of the activation of the sensing device was assessed by performing continuous culture of the autocrine strains in eVOLVER bioreactors^41^. The ReacSight strategy was used to connect these bioreactors to a flow cytometer through a pipetting robot for automated monitoring of cell fluorescence^42^. For both displayed and secreted mNeonGreen (dNL and sNL), there is an overshoot in downstream activation after light induction, followed by stabilization to steady state levels. Therefore, the downstream activation produced by the device is stable over time.

### Growth-based biosensor for improved secretion or display

After demonstrating that our device produces an autocrine-only signal that scales with POI display/secretion and is stable over time, we investigated whether this would allow us to control growth rate in selective environments.

The lack of a nutrient in the environment and the corresponding prototrophy are widely used in yeast to select for a desirable genotype^39,43^. In our *his3Δ0* chassis, tuning *HIS3* expression allows control of the growth rate in medium lacking histidine (Figure 1E). Addition of 3-aminotriazole (3AT) in his-medium demands higher *HIS3* expression^39,44^, allowing higher stringency that expands the range of downstream activation for which growth rate can be tuned quantitatively. We sought to determine the relationship between *HIS3* expression, medium stringency and growth rate using graded optogenetic induction of mScarletI.2a.*HIS3* (Figure 3A, Supplementary Note 4). mScarletI fluorescence acts as a proxy for *HIS3* expression. As expected, induction with varying duty cycles produced graded levels of reporter expression. In his+ medium, growth rate remained above 0.35 h^−1^, regardless of reporter expression. As shown in Figure 1E, a range of growth rates is observed in his-medium, with 40% DC being sufficient to recover a growth rate above 0.35 h^−1^. The addition of 3AT reduced the overall growth rate, while preserving the gradation across induction levels. Plotting growth rate as a function of reporter expression illustrates how medium stringency modulates the relationship between the two. In his-medium, a graded relationship is observed between 0 and 40 mScarletI units of fluorescence, a range that corresponds to the downstream activation levels observed in the autocrine strains sNL and dNL (Figure 2B). Therefore, using the mScarletI.2a.*HIS3* reporter in autocrine strains should quantitatively link growth rate and secretion in his-medium.

**Figure 3:**
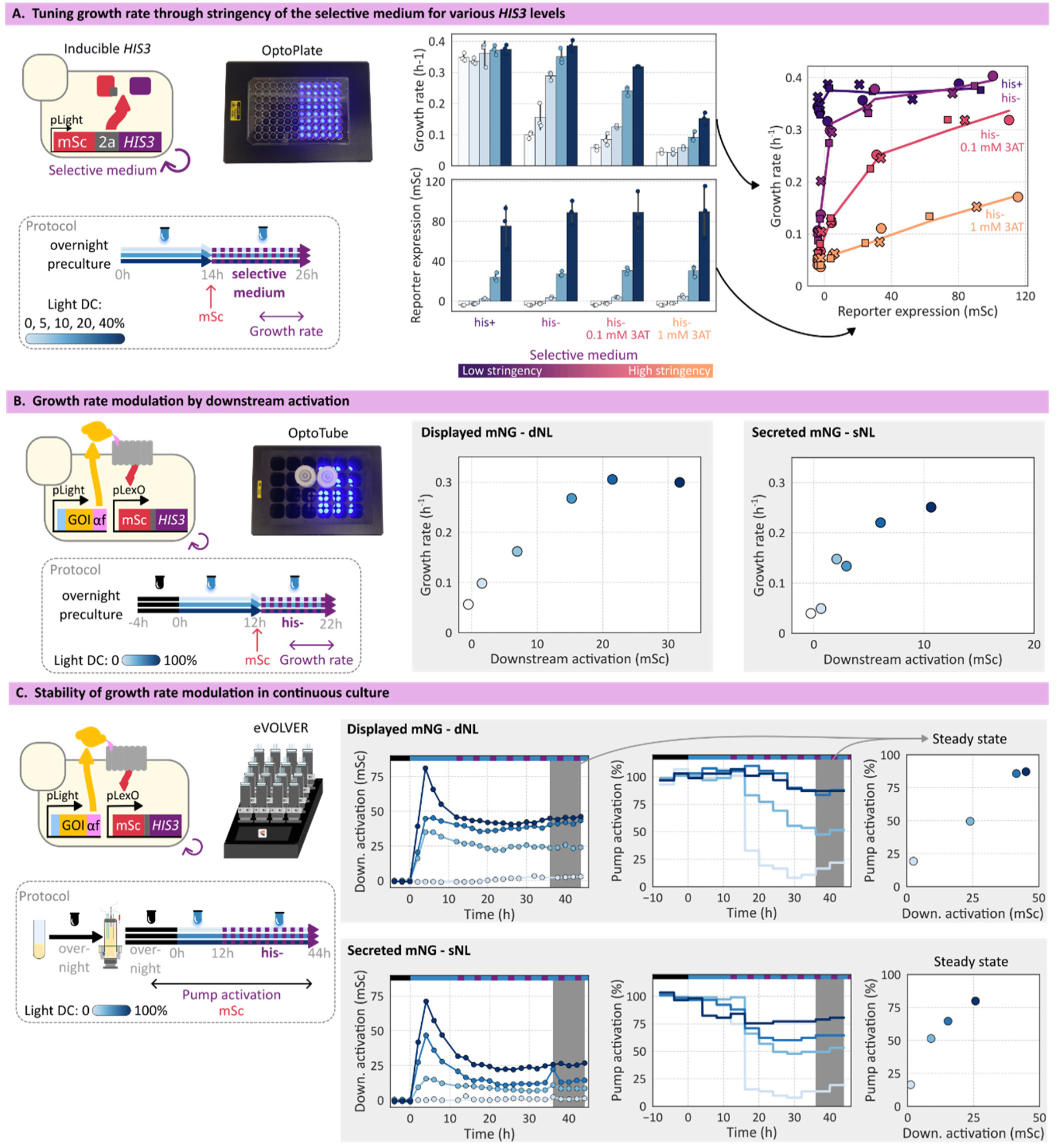
Coupling growth to secretion through histidine prototrophy recovery. **A)** Assessment of *HIS3*-dependent growth under varying stringency. Cells expressing mScarletI.2a.*HIS3* reporter from pLight promoter are grown under blue light in his+ medium and their fluorescence measured by flow cytometry before being transferred into fresh medium of increasing stringency (his+, his-, his-0.1 mM 3AT and his-1 mM 3AT). Multiple OD measurements are taken during the 2^nd^ incubation to determine growth rates. Experiments are done in biological triplicate; medians of each population are represented by dots, and their mean and corresponding standard deviation are shown with the bar plot. **B)** Assessment of mNG display/secretion-dependent growth in batch. Autocrine strains, displaying or secreting mNG, containing the mScarletI.2a.*HIS3* reporter are grown under blue light in his+ medium before cell fluorescence measurement by flow cytometry and transfer into fresh his-medium. Multiple OD measurements are taken during the 2^nd^ phase of the culture to compute growth rates. Median fluorescence of each population is shown. **C)** Assessment of mNG display/secretion-dependent growth in continuous culture. Autocrine strains are grown in turbidostat in 40% histidine medium in the dark to allow cells to reach exponential growth. Then, various blue light duty cycles are applied and after 12 hours of culture the medium is switched to his-. Cell fluorescence is measured every 2 hours by automated flow cytometry and median fluorescence of each population is shown. Thanks to the turbidostat regime, pump activation, which represents dilution rate, is used as a proxy for the growth rate. Steady states values are obtained by taking the mean from the last 8 hours of culture.

We built the corresponding autocrine strains and assessed their growth rate in his-medium under varying light duty cycles, using a 2-phase culture to first induce POI expression before placing cells in selective medium (Figure 3B, Supplementary Note 4). As expected, growth rates scaled with downstream activation for both displayed and secreted mNG, confirming that his-stringency was appropriate to generate differences in growth rate. We demonstrated previously the monotonic relationship between secretion/display and downstream activation. Taking these results together with the absence of paracrine activation suggests that our biosensor is able to translate improved secretion/display into faster growth at the single-cell level, thereby allowing growth-based screening of pooled libraries.

Growth-based screening is most effective in continuous culture, where cells are maintained in exponential growth throughout the selection process, as opposed to batch culture in which lag and stationary phases are inoperative. Hence, we decided to test whether the previous results held in continuous cultures (Figure 3C). The same autocrine strains were subjected to a range of light duty cycles and grown in a medium containing 8 mg/L histidine, representing 40% of the usual concentration, before switching to his-medium. Activation of the input pump was used as proxy to track growth rate, since dilution rate and growth rate are equal in these conditions (Supplementary Note 4). As observed in batch culture, graded secretion or display of mNG led to graded downstream activation, which itself generated graded growth rates upon his-medium switch. Importantly, growth rates are stable at steady-state, except for 0% duty cycle showing a small but limited regrowth. These results suggest that the AutoGrowth strategy can be used to screen pooled libraries for improved secretion in continuous cultures.

### Generalization of the biosensor to other POIs

After demonstrating the system properties with the easy-to-secrete mNG protein, it appeared essential to assess the biosensor performance with POIs of biotechnological and medical interest (Supplementary Note 1). We included two xylanases (XylC and Xyl2), an α-amylase (Amy), a nanobody (Nb21), and a single-chain variable fragment (scFv).

Autocrine strains were built to secrete such POIs fused to mNG and α-factor (Figure 4A). Graded induction of POI secretion was applied to investigate the relationship between secretion and downstream activation. We used a 2-phase culture, as in Figure 3B, to focus on steady-state behaviors, which are more relevant for later applications, that is, screening in continuous culture. All secreted POIs generated a clearly detectable signal. For all POIs but Nb21, a monotonic relationship was observed between secretion and downstream activation, supporting the ability of the biosensor to screen for improved secretion of these POIs. For Nb21, one can see that a strong induction resulted in high downstream activation, low secretion levels, and significant fitness cost. Therefore, despite having a high activation level, it is unlikely that these poor secretors would be selected during growth-based screens. We also observed that POIs that are highly secreted tended to cause lower downstream activation (Supplementary Note 5). We hypothesized that efficiently secreted POIs navigate efficiently across the secretory pathway and the cell wall, while hard-to-secrete POIs, especially scFv and Amy, accumulate in the secretory pathway or in the periplasm, and have more time to activate Ste2id. To evaluate this hypothesis, we first explored the subcellular localization of these POIs (Figure 4B, Supplementary Note 5). They all showed vacuolar accumulation, while for few cells, there was cytoplasmic accumulation. Importantly, a signal at the cell periphery was observed only for cells secreting scFv and Amy, especially for the latter (red arrows), suggesting that higher downstream activation comes from periplasmic accumulation. To confirm it, we tested the effect of the addition of dTA, the α-factor antagonist already used in Supplementary Note 3, on the activation caused by secreted scFv and Amy (Figure 4C). The rationale is that exogenously added dTA precludes the activation, by secreted scFv or Amy, of receptors present at the cell surface. As expected, dTA addition caused major reduction of the downstream activation, confirming that secreted scFv and Amy activate Ste2id at the cell surface. Hence, downstream activation is a proxy for the periplasmic concentration of proteins. Consequently, genetic variations increasing secretion through improved cell wall diffusion would be missed by the biosensor. Nonetheless, any modification leading to higher concentrations in the periplasm would be detected.

**Figure 4:**
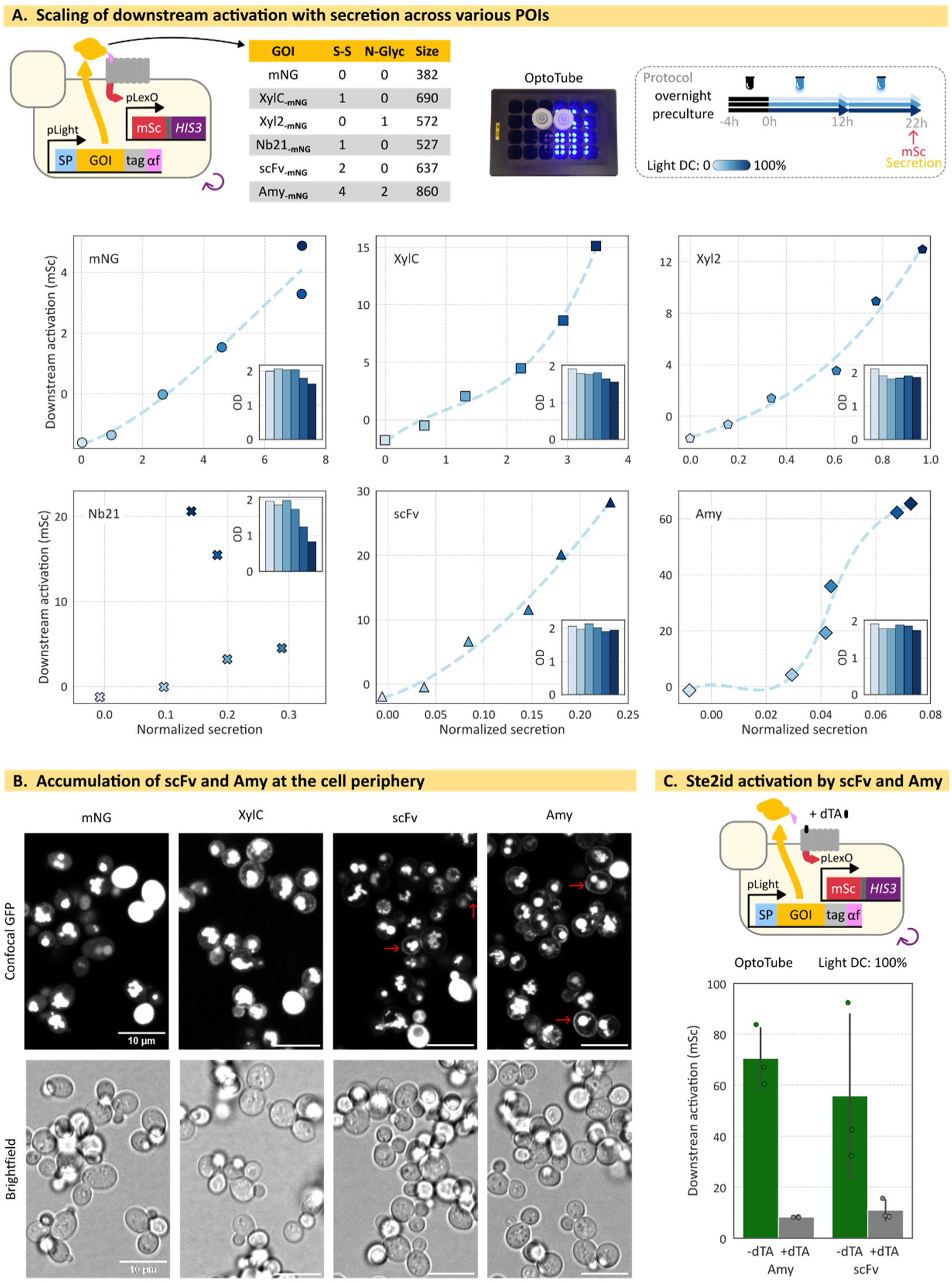
AutoGrowth biosensor to screen various POIs. **A)** Assessment of the relationship between secretion and downstream activation. Autocrine strains secreting various POIs are grown in his+ medium under a range of blue light for 12h before inoculation of fresh his+ medium for 10 hours before measurement by flow cytometry of cell fluorescence and secretion, through anti-Flag magnetic beads. Median fluorescence of each population is shown. **B)** Surface accumulation pattern is protein specific. Autocrine strains are grown in his+ medium under 100% blue light DC for 12 hours before inoculation of fresh his+ medium for 10 hours and transfer of samples into wells of ibidi µ-slide for confocal microscopy (60x). **C)** POIs at the cell surface trigger downstream activation. Autocrine strains secreting scFv or Amy are grown for 12h under light induction in his+ medium supplemented or not with dTA at 5 µM final concentration, before transferring into fresh his+ medium supplemented with dTA at 5 µM final concentration for 10 hours before measurement by flow cytometry of cell fluorescence. Experiments are done in biological triplicate; medians of each population are represented by dots, and their mean and the corresponding standard deviation is shown with the bar plot.

### Demonstration of AutoGrowth capacity to screen pooled libraries and select rare favorable variants

Finally, we investigated whether the biosensor can be used to screen pooled libraries and enrich rare, favorable variants. scFv was chosen as POI for these proof-of-concepts because microbial production of antibody fragments is promising but challenging, thereby often requiring strain optimization. We first confirmed the biosensor ability to couple growth to scFv induction in continuous culture under mild stringency (Supplementary Note 6).

We chose to screen for the best signal peptide (SP), an essential component of secretion optimization. Previous strains in this study were built with MFo2, an optimized version of the pre-pro-peptide from α-factor^45^. We selected 12 new SPs studied previously^46–48^ and built the corresponding autocrine strains secreting scFv (Figure 5A top, Supplementary Note 7). The 13 strains were first assessed in batch monocultures with OptoTube to establish the “ground truth”, that is, the scFv secretion and downstream activation, for each SP. We rank them based on the secretion levels. No OD-normalization is performed here, since we aim at screening for the SP giving the best secretion, irrespective of the growth. We observed various secretion levels, validating the relevance of the SP library to test the biosensor. Unsurprisingly, final OD tended to be lower in strains with stronger secretion. This fitness defect reveals a burden generated by strong secretion. Importantly, downstream activation scaled with secretion, and efficient SPs generated the strongest signals, suggesting that the biosensor can be used to screen for the best SPs. Confocal microscopy showed that subcellular localization of scFv is strongly dependent on SP efficiency, with cytoplasmic accumulation for inefficient SPs and periplasmic accumulation for all efficient SPs except MF8 (Supplementary Note 7).

**Figure 5:**
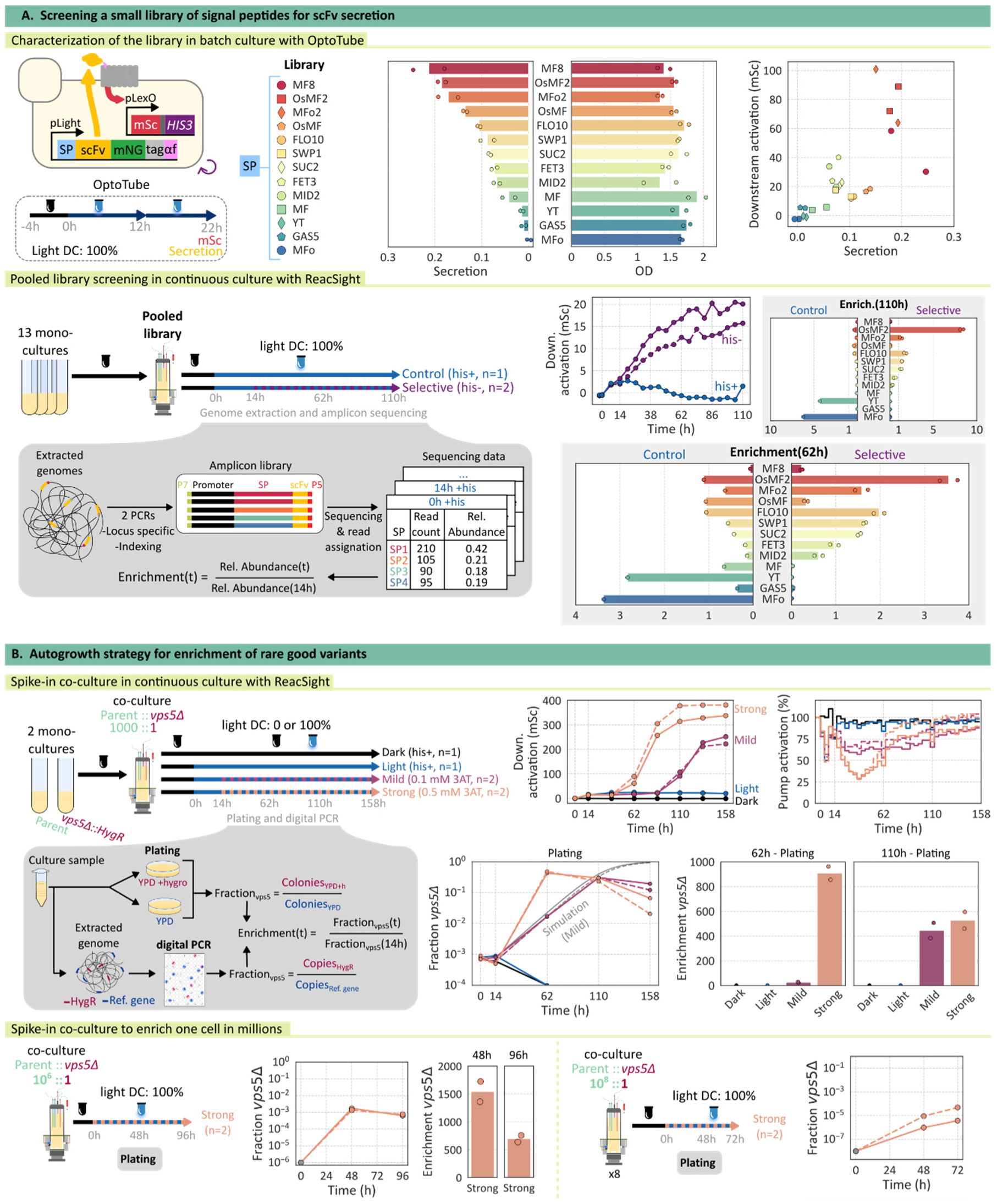
AutoGrowth biosensor to screen pooled libraries and enrich scarce variants. **A top)** Assessment of the biosensor for scFv secretion with several SPs. Autocrine strains secreting scFv with various SPs are grown in OptoTube setup in his+ medium for 4 hours in the dark then for 12 hours under 100% blue light DC, before inoculation of fresh his+ medium for 10 hours under blue light followed by flow cytometry measurement of cell fluorescence and secretion, through anti-Flag magnetic beads. Experiments are done in biological duplicates; medians of the population are represented with dots and their means shown by the bar plot. **A bottom)** Assessment of small-scale pooled library screening. Autocrine strains secreting scFv with various SPs are mixed in equimolar ratio and grown in eVOLVER bioreactor with turbidostat in 40% histidine medium in the dark before application of 100% blue light DC and later switch to his-, except for controls. Cell fluorescence is measured every 6 hours by automated flow cytometry, with medians of distributions being represented. Samples taken at the specified timepoints are subjected to genome extraction, amplification of the SPs and sequencing with short-read Illumina sequencing (MiSeq100, 5 million reads flow cell). Reads are mapped to SPs and their count normalized by the total number of reads per sample, defining the relative abundance used to compute enrichment. **B top)** Assessment of the biosensor for enrichment of good variants among thousands of competitors. The *vps5Δ* and parental strains are mixed in ratio 1:10^3^ and grown in eVOLVER bioreactor in turbidostat in 40% histidine medium at first in the dark before application of 0 or 100% blue light DC for 14 hours before switch, when applicable, to his-medium supplemented with 0.1 or 0.5 mM 3AT. Cell fluorescence is measured at light induction, medium switch and then every 24 hours by automated flow cytometry, with medians of the population of each sample being represented. Samples are taken at the specified timepoints, diluted if needed, and plated onto YPD agar medium supplemented or not with hygromycin because the *vps5Δ* strain contains a hygromycin resistant cassette but not the parental. Colonies are counted to then compute the fraction of the *vps5Δ* cells throughout the course of the culture. **B bottom)** Assessment of the biosensor for enrichment of good variants among millions of competitors. The parental and *vps5Δ* strains are mixed in ratio 1:10^6^ or 1:10^8^, and grown in eVOLVER bioreactor in turbidostat in 40% histidine medium at first in the dark before application of 100% blue light DC and medium switch to his-medium supplemented with 0.5 mM 3AT. Samples are taken at the specified timepoints, diluted or concentrated if needed, and plated onto YPD agar medium supplemented or not with hygromycin because the *vps5Δ* strain contains a hygromycin resistant cassette but not the parental. Colonies are counted to then compute the fraction of the *vps5Δ* cells throughout the course of the culture.

We then set out to screen the pooled SP library of the 13 strains in eVOLVER bioreactors (Figure 5A, bottom). The library was grown in continuous culture, either with or without selection, to compare the enrichment allowed by our biosensor with the natural drift based solely on fitness. We observed that median downstream activation increased through the culture only in selective medium, advocating for the successful selection of cells with higher downstream activation. Cultures were sampled at light induction (0h), selection start (14h), and after 2 or 4 days of culture (62h and 110h). The genomic region containing the SP was PCR amplified and sequenced using short-read technology to compute strain enrichment. We observed, in absence of selection, that the proportions drifted away from the initial state at 14h, reflecting the effect of SPs on the burden associated with POI secretion. MF8, FED3, and MID2 were clearly burdensome, because they were depleted after only 2 days. By contrast, YT and MFo were strongly enriched, indicating that scFv production was the least costly for these 2 strains. In selective conditions, and after 2 days of culture (62h), SP enrichment mirrored secretion efficiency for most variants, except OsMF and MF8. For the latter, it likely reflects the altered fitness already observed without selection, confirming the ability of AutoGrowth to account for fitness of variants during screening. Importantly, one of the most efficient SP in batch culture, OsMF2, was strongly enriched after 2 days, and even more after 4 days. These results demonstrate that the biosensor can be used to screen pooled libraries for both improved secretion and robust fitness in lab-scale bioreactors.

The enrichment ratio obtained in the SP screening indicates that best variants could have been found by plating the population and testing few clones individually. However, this is strongly dependent on the library size and could not be feasible for variants present in very low abundance at the culture onset. We sought to test the capacity of the biosensor to enrich good variants present in low or very low abundance. First, we assessed several genetic perturbations with the biosensor and found that the deletion of *VPS5* increases scFv secretion 4-fold and downstream activation 6-fold (Supplementary Note 8). The beneficial effect of this deletion was confirmed in BY4741 background free from mating pathway engineering (Supplementary Note 9). The autocrine v*ps5Δ* strain also outperformed the parental strain in bioreactor under mild stringency (his-0.1 mM 3AT), with growth rates of 0.29 h^−1^ against 0.22 h^−1^. Using these growth rates, the simulation of a co-culture of these two strains showed that, starting from 0.1%, *vps5Δ* cells could reach 40% of the population in 4 days (Supplementary Note 9). We performed the corresponding spike-in experiment, added two control experiments (absence of selection, with or without light induction), and a more stringent condition (his-0.5 mM 3AT). Results are shown in Fig 5B. In the absence of selection, population-averaged downstream activation and growth remained constant, irrespective of the light induction, highlighting the stability of the system. Abundance of the *vps5Δ* strain, that harbors an hygromycin resistance cassette, was followed by sampling the co-culture and plating cells on YPD with or without hygromycin. The *vps5Δ* fraction decreased through the culture for both controls, indicating an altered fitness of the *vps5Δ* strain independent of scFv production that was not observed in batch (Supplementary Note 8). Another explanation could come from a potential burden caused by the hygromycin resistance cassette, absent in the strain used for the batch culture. In the presence of selection, population-averaged downstream activation started to increase after 2 and 4 days under strong and mild stringency, respectively, indicating enrichment of cells with high activation. Growth of the population slowed down transiently after the start of selection, with strong stringency causing a stronger, yet shorter, growth decay. Under mild selection, the *vps5Δ* fraction was 0.07% at the start of selection (14h), and reached 30% after 4 days of culture, representing a 400-fold enrichment. Data and simulation agree on days 2 and 4, but not 6, because of a slight decrease of *vps5Δ* fraction. Under strong selection, the *vps5Δ* strain represented 0.06% at the selection start and exceeded 40% of the culture in only 2 days, representing an 800-fold enrichment. The *vps5Δ* fraction was also assessed by digital PCR, and results agreed with those obtained using the plating method (Supplementary Note 10). These results demonstrate the power of the AutoGrowth biosensor to enrich rare variants in limited time. We investigated further the slight decrease of *vps5Δ* fraction after it peaked and found that a “cheater” behavior appears (Supplementary Note 11). Cheater cells show high downstream activation decoupled from their secretion capacity. They recover normal growth, regardless of the selection stringency. When this happens, fitness becomes the only driver of selection, which allows cheater cells to outgrow *vps5Δ* cells, as observed in the control samples. Consequently, high stringency should be applied for a limited duration to avoid appearance of cheaters. Notably, AutoGrowth was >10 times more efficient to enrich *vps5Δ* cells among the co-culture than one round of FACS, taking the top 0.3% cells with the highest downstream activation signal (Supplementary Note 10).

The throughput of FACS or FADS is determined by the number of cells or droplets that can be screened. By contrast, the screening throughput of the AutoGrowth strategy is limited by the size of the culture and the capacity to enrich good mutants before emergence of cheaters. We sought to challenge the capacity of our biosensor and conducted a spike-in experiment with a *vps5Δ*:parental ratio of 1:10^6^. After 2 days under strong stringency, *vps5Δ* cells were enriched 1500-fold and accounted for 0.15% of the population. Since large library in yeast can reach up to 10^8^ variants^34^, we pushed further our tests and assessed a *vps5Δ*:parental ratio of 1:10^8^. Because of the very large dilution factor, we used 8 bioreactors together to have an estimate of 10 *vps5Δ* cells in total at culture onset. After 3 days, the fractions of *vps5Δ* were higher than 10^−6^, with an average enrichment higher than 2500. This is high enough to be easily retrieved by deep sequencing. Such growth-based strategies can therefore outperform the throughput of snapshot-based methods for library screening.

## Discussion

We presented AutoGrowth, a strategy allowing growth-based screening for improved secretion in yeast, using a biosensor coupling growth to secretion at the single-cell level. We first demonstrated the strict single-cell nature of the biosensor and its ability to modulate growth proportionally to the secretion or display levels of mNeonGreen. The genericity of the sensor was then proven with 5 POIs of industrial or biomedical relevance, which present various secretion challenges and span a range of secretion levels. Finally, we performed proof-of-concept experiments to test AutoGrowth capacities in pooled conditions. First, we screened a small-scale library of signal peptides for scFv secretion by coupling AutoGrowth with deep sequencing. Variants showing high secretion and robust fitness were successfully enriched. Second, we explored the capacity of the biosensor to screen large-scale libraries via spike-in experiments. We were able to identify good variants in one-to-one-million and in one-to-100-million competitions, with thousand-fold enrichments in a few days. Such enrichments would be easily detected by deep sequencing, demonstrating the capacity of the biosensor to screen barcoded libraries containing hundreds of millions of variants.

In the AutoGrowth strategy, we measure secretion capacity through the enrichment/depletion of a variant. This readout results from the temporal integration of growth rate differences between variants. There is therefore an exponential increase of the enrichment/depletion ratios with increasing selection times. This distinguishes our method from those using snapshot measurements (*i.e.* FACS and FADS) to sort variants. Another key feature of AutoGrowth is the possibility to screen for either the sole secretion signal or a balance of secretion and fitness by adjusting the stringency of the medium. Selection under strong stringency overrides any fitness defect coming from secretion burden or genetic perturbations, making secretion capacity the dominant driver of selection, whereas mild stringency favors variants with optimal secretion/fitness balance. Such tunability is useful to adapt the screen to the needs of different production modes (batch, fed-batch, and continuous culture). Of note, strong stringency accelerates the selection of good variants, yet it also causes the appearance of cheater cells. These cheater cells upregulate the expression of the selection marker, decoupling it from secretion capacity, which results in the selection operating on fitness only, and not on the combination of secretion and fitness. This phenomenon appears after at least 2 days of culture in our conditions, leaving enough time for substantial enrichment before. Adding counter selection steps inhibiting cells whose reporter expression is decoupled from POI secretion could alleviate this limitation. We could leverage systems based on *CAN1* and *FUI* that have already been used with the yeast mating pathway^49,50^.

Many methods have been developed to screen for improved secretion in yeast. Approaches based on 96-well plates offer simple screening methods^10–13,15^, particularly useful to screen transformants in the presence of clonal variation. However, they fail to screen libraries with thousands of clones or require heavy automated platforms. Microdroplet-based approaches preserve genotype-phenotype association thanks to single cell encapsulation and allow higher throughput^9,16–22^. The screening of various libraries (CRISPRi/a, RNAi, and UV-mutagenesis) led to the identification of several beneficial genetic perturbations generating great insight into secretion optimization. Most experiments relied on the enzymatic activity of α-amylase, but few POI-agnostic approaches were also developed^20,21^. The relevance of these microdroplet-based approaches is limited by the fact that the micro-environment of cells is strictly constrained and significantly different than liquid fermentation conditions^23,24^. Consequently, variants identified in microfluidic screens could be specific to cell state in micro-environment. This could explain why, in most studies, less than 60% of variants from the sorted population show secretion higher than the parental strain^17–19,21,22^. More importantly, fermentation-specific beneficial perturbations could be missed. Other methods screen pooled libraries on a solid medium, with limited diffusion of the POI, resulting in good colonies showing a noticeable phenotype^8,25–27^. These approaches combine relative high throughput with simplicity. However, they are often based on a specific enzymatic reaction making the assay POI-specific or requiring fusion of the POI with the corresponding enzyme. Of interest, Cleaver *et al.* recently proposed a generic approach that is based on co-secretion of the POI and α-factor, the latter being sensed by Ste2 to produce a fluorescent signal^8^. Their biosensor shares many similarities with our system. However, the relevance of their sensing module is limited by its very high sensitivity, due to the use of the strong promoter pCCW12, and by the potential intracellular inactivation of the wild type Ste2^37^. In addition, their system is likely to be non-functional in liquid conditions, since we found that α-factor fusion is required to have a strict single-cell sensor. It constrains their screening to be performed on solid medium. Yet, cells growing on solid medium have different physiological states than cells growing on liquid cultures^28^. Methods based on the capture-and-secrete approach suffer from similar limitations, since they need tailored screening conditions, such as high viscosity or static liquid culturing^29,30^. Methods based on Yeast-Surface Display (YSD) make it possible to screen in liquid culture because the POI is captured on the cell wall thanks to its fusion to an anchor. Display is measured at the single-cell level with antibody labeling of cells and FACS, permitting the screening of pooled libraries for improved display. It has been successfully applied to screen a genome-wide library of cDNA and a library of 80 promoter-SP combinations^33,32^. YSD is rather versatile since it does not necessitate an engineered chassis, the only requirement being Aga1 overexpression to efficiently capture the Aga2-fused POI. Yet, several technical features make the connection between the display signal, measured through quantifying immuno-labeled anchored proteins, and the actual secreted levels not always straightforward. One can mention the interference of the Aga2 anchor, which forced Wentz *et al.* to restrict their screen to their most complicated POI under restrictive temperature^33^. The presence of an affinity tag, used for immuno-labeling, between the signal peptide and the POI could also interfere, since residues around the cleavage site of the SP were shown to be involved in secretion efficiency^51^. In addition, when applied to large libraries (10^6^ variants or more), the capacity of cytometry snapshot measurements to not miss rare, good secretors using immuno-labeled anchored proteins is not clearly established. For instance, in their Gigantic Parallel Reporter Assays, De Boer *et al.* constructed a library of 10^8^ random DNA sequences to study transcription regulation, but less than 10^8^ cells were sorted through FACS, preventing evaluation of all variants^34^. By contrast, we demonstrated the capacity of our growth-based single-cell biosensor to identify variants of interest in such large libraries. In fact, the throughput of our method is only limited by the sequencing capacity, which is about 10^10^ reads for modern production-scale sequencers. In summary, our method based on an autocrine growth-based sensor stands out as the only method allowing to screen pooled libraries for improved secretion of a POI in liquid culture.

The secretion capacity of cells depends on many diverse parameters, which makes optimizing secretion challenging. Current models cover only a fraction of the parameter space, such as signal peptide^52–54^, 5’UTR/promoter^34,55,56^, and codon-usage^57,58^. Technologies like Golden Gate cloning and CRISPRa/i facilitate the construction of large-scale libraries, which interrogate further the parameter space. By enabling the screening of very large libraries at minimal resource and equipment cost, AutoGrowth could play a pivotal role in generating the comprehensive datasets required to develop predictive sequence-to-function models using machine learning. Such models are expected to play a key role in protein engineering, directed evolution, and synthetic biology.

## Methods

### Cloning

*Escherichia coli* strain XL1-Blue (Agilent Ref. 200249) was used for cloning, it was grown at 37°C in Lysogeny Broth (LB) medium. Plasmids used in this study are listed in the Supplementary Note 1. They have been constructed using the framework of the Yeast Tool Kit, a widely used Modular Cloning (MoClo) kit^59^. Plasmids from other compliant kits were also used: Yeast GPCR-sensor Toolkit (Addgene Ref. 1000000157)^35^, Multiplex Yeast Toolkit (Addgene Ref. 1000000229)^60^ and EasyClone-MarkerFree Vector Set (Addgene Ref. 1000000098)^61^. Additionally, new genetic parts were added to the MoClo system by amplification from the genome of *S. cerevisiae* BY4741, or purchased from Twist Biosciences, IDT and Eurofins depending on specific needs. Golden Gate reactions were performed in a volume of 10 µL with 1X T4 ligase buffer, BSA (Thermo Scientific Ref. B14), restriction enzyme (NEB, Ref. R3733L or R0739L) and T4 DNA ligase (Promega Ref. M1804). Level 0 plasmids were systematically sequenced upon storage. Plasmids used in the signal peptide library were constructed with an automated Golden Gate workflow from the SlowPoke framework^62^, using Golden Gate kit (NEB Ref. E1601L) with a 12 µl final volume.

### Yeast strains and culture conditions

Listed in the Supplementary Note 1, yeast background strains used in this study are BY4741 or yWS677, which itself originates from BY4741^35^. Genetic modifications were performed with the lithium acetate/single-stranded carrier DNA/PEG transformation method from Gietz and colleagues^63^. Genetic materials were either episomal plasmids or linear fragments (digested plasmids or PCR products) bearing an auxotrophic or an antibiotic resistance marker. For marker-less integration and genome editing, CRISPR-Cas9 based transformations were performed. In such cases, the integrative linear fragment (digested plasmids, PCR products or annealed 120bp oligonucleotides purchased from Eurofins) was co-transformed with a 2µ episomal plasmid expressing Cas9, the sgRNA and the G418 resistance marker. Correct integration was verified by junction colony PCR (DreamTaq, Thermo Scientific Ref. K1081), followed by genomic DNA extraction (Biosearch Technologies Ref. MPY80200), high-fidelity amplification of the engineered locus (Phusion, Thermo Scientific Ref. F530L), and Sanger or ONT amplicon sequencing (Eurofins). Transcription factors NLS-VP16-EL222^64^, for optogenetic induction, and LexA-PRD_STE12_^35^, for downstream activation, were expressed from the *URA3* locus, with promoters pTDH3 and pRAD27 respectively. The sensing module made of *GPA1* and *STE2*/*STE2id* was expressed from the XI-2 locus^61^ with the promoters pPGK1 and pCCW12/pPOP6. The reporter, mScarletI or mScarletI.2A.*HIS3*, was expressed from the *HO* locus with the LexO6x.pLEU2m promoter^35^.

The POI production module was expressed from the *LEU2* locus with pLight promoter, originally named 5xBS-CYC180pr^65^. As described in Figure 1B, the POI module contains several domains from N to C-terminal: a signal peptide, the gene encoding the protein of interest, Aga2 in the case of displayed proteins, a FLAG-HIS tag and the α-factor. GS linkers are added between domains of the fusion protein. For gene deletion, the entire ORF was removed or replaced with a hygromycin resistance cassette. For the overexpression of endogenous genes, the corresponding gene and rtTA-Gal4^66^ were expressed from the Int-7 locus^60^ with the promoters pTET and pRNR2 respectively.

Yeasts were grown at 30°C with shaking in YPD or in SC LoFlo medium (1X YNB - ForMedium Ref. CYN6505, 2 g/L glucose - Sigma G7021, 1X CSM - ForMedium Ref. DCS0019). For experiments in OptoPlate, OptoTube and eVOLVER bioreactors, cells taken from −80°C glycerol stock were streaked on YPD plates for 2 days at 30°C. Precultures were prepared in SC LoFlo medium inoculated with single colonies.

### Batch optogenetic experiments in tubes - OptoTube

Overnight precultures were used to inoculate 2 mL of SC LoFlo supplemented with 25 mM phosphate buffer (1M stock with 125.41 g/L Sigma-Aldrich Ref. S0751 and 23.14 g/L Sigma-Aldrich Ref. P8281; buffered LoFlo) to OD_600_ 0.05, and grown for 4 hours in the dark then 10 hours under blue light illumination with the OptoWELL 96 device (Opto Biolabs GmbH) using a specific adaptor for tubes. Experiments were conducted at 30°C with shaking at 500 rpm. For experiments with two phases under light induction, 2 mL of buffered SC LoFlo were inoculated from overnight culture to OD_600_ 0.025 for 4 hours of growth in the dark and 12h with blue light induction, at 30°C shaken at 500 rpm. Then, pre-warmed 2 mL of buffered SC LoFlo were inoculated with induced cultures to OD600 0.1 and grown for 10 hours. Following the 10-hour incubation in light, the OD_700_ was measured with the plate reader (Tecan Spark). In addition, cell fluorescence, secretion and display were measured by flow cytometry (Cytek Guava easyCyte BGV HT). When applicable cells were imaged with confocal microscopy (Oxford Instruments BC43).

For experiments in selective conditions, buffered SC LoFlo lacking histidine (1X CSM-His, ForMedium Ref. DCS0079) was used at the resuspension step (12h post light induction) supplemented when needed with 3-aminotriazole (3AT, Sigma-Aldrich Ref. A8056). A 3AT stock solution was prepared fresh each time at a concentration of 20 mM.

For experiments with desTrp1,Ala3 α-factor (dTA, Genscript), protocols were identical except that dTA (or a corresponding water volume) was added 1 hour prior to light induction at 5 µM final concentration.

When needed, doxycycline (dox, Sigma-Aldrich Ref. D5207) was added in the buffered SC LoFlo medium in which overnight cultures were resuspended as well as at the resuspension step 12h after light induction. Dox was used at a final concentration of 1 µg/mL.

### Batch optogenetic experiments in 96-well plate - OptoPlate

Precultures were used to inoculate 200 µL of fresh SC LoFlo at OD_600_ 0.05 in wells of a microplate (Greiner bio-one Ref. 655090). Fresh culture was grown for 14 hours at 30°C and shaking at 700 rpm, with *HIS3* induction through light illumination with the OptoWELL device. Cell fluorescence was assessed by flow cytometry (Cytek Guava easyCyte BGV HT), then 5 µL of culture were used to inoculate pre-warmed SC LoFlo medium with or without histidine and 3AT for a total volume of 200 µL. OD_700_ was followed with measurements in the plate reader (Tecan Spark).

### eVOLVER bioreactor experiment with the ReacSight platform

Precultures were used to inoculate eVOLVER vessels (FynchBio) filled with 25 mL SC LoFlo medium buffered with 3.75 mM of arginine. For experiments using selective medium, the starting medium contained 40% of SC LoFlo and 60% of histidine lacking SC LoFlo, resulting in 8 mg/L of histidine instead of 20 mg/L. For the library and spike-in, OD_700_ of the precultures was measured to mix strains at the desired ratio prior to inoculation. A recovery period lasting at least 8h in the vessels was performed before the start of automated flow cytometry measurement (Cytek Guava easyCyte BGV HT). Samples are processed automatically with the pipetting robot OT-2 (Opentrons). Depending on experiments, these measurements were performed every 2, 6 or 24 hours. The target OD of the turbidostat regime was set to 0.5, temperature to 30°C and stirring to 8. Light induction was performed thanks to the addition of a blue LED to each eVOLVER bioreactor.

For experiments requiring sampling of the culture for plating and genome extraction, 3-5 mL of cultures were sampled using the sampling lines of the ReacSight platform. For amplicon sequencing and digital PCR, 2.8 mL of culture were used for genome extraction (Biosearch Technologies Ref. MPY80200), and the concentrations of resulting samples were measured with a Qubit 2.0 Fluorometer (Invitrogen). For plating, 10-fold dilutions were made in a 96-well plate before transferring 50 µL of the relevant dilutions onto YPD supplemented or not with hygromycin at 600 μg/mL. Colonies were counted using a counting pen (Thermo Scientific Ref. 11862710).

### α-factor dose-response curve

Precultures were used to inoculate 2 mL of buffered SC LoFlo at OD600 0.05, and grown for 4 hours at 30°C and 500rpm. α-factor (Sigma-Aldrich Ref. 63591) was added at the indicated concentrations, and cells were incubated 10 hours at 30°C and 500 rpm. Cell fluorescence was assessed by flow cytometry. When needed, dTA (5 µM final concentration) or a corresponding water volume was added 1 hour prior to addition of α-factor.

### Secretion measurement

Secretion measurements were adapted from a previously established protocol^40^. Briefly, following OptoTube experiments, 50 µL of culture were mixed with 150 µL of sterile water, 26 µL of Cerulean sample, 26 µL of 100 mM phosphate buffer, and 10 µL of washed anti-Flag magnetic beads (Thermo Scientific Ref. A36797). The solution was mixed by pipetting and incubated at room temperature, in the dark for 1 hour. Samples were then washed 3 times with 200 µL of TBS, using a magnetic rack to immobilize the beads. Finally, samples were resuspended into 200 µL of sterile water and their fluorescence measured by flow cytometry (Cytek Guava easyCyte BGV HT). Cerulean samples were prepared in house by cultivating a strain secreting Cerulean and freezing the culture supernatant at - 80°C for later use as internal control allowing normalization during data processing. Anti-Flag beads were prepared before each experiment as follows: 0.4 µl of vendor solution per sample were washed twice with TBS and resuspended in sterile water before their use, 10 µL per sample.

### Antibody-based display measurement

Following OptoTube experiments, 10^6^ cells were sampled and washed three times with 200 µL of cold PBS-BSA buffer (PBS with 1% BSA at pH 7.5), with centrifugation step of 3 minutes at 6000 g. Cell pellet was resuspended in 100 µL of PBS-BSA buffer supplemented with 1:200 dilution of primary anti-FLAG M2 antibody (Sigma-Aldrich Ref. F3165-.2MG) and incubated 90 minutes at room temperature with gentle agitation. Cells were then washed three times with 200 µL of cold PBS-BSA buffer, and resuspended in 100 µL of cold PBS-BSA buffer supplemented with 1:200 dilution of secondary goat anti-mouse antibody coupled to Alexa Fluor™ Plus 405 (blue fluorescent dye, Invitrogen, Ref. A55057) and incubated 90 minutes at room temperature with gentle agitation. Finally, cells were washed three times with cold PBS-BSA buffer, resuspended in 200 µL of PBS pH 7.5 and sent for flow cytometry (Cytek Guava easyCyte BGV HT) or confocal microscopy (Oxford Instruments BC43).

### Flow cytometry data analysis

For cell fluorescence measurement by flow cytometry, we applied a previously developed^42^ automated procedure for gating of single cell and deconvolution of fluorescent signal (Supplementary Note 12). When needed, strains contained in co-culture were identified thanks to their blue fluorescence using automated clustering with a Gaussian mixture, as shown in Figure 2A.

For display, cell fluorescence was measured by flow cytometry and processed as presented above. The BLU-V values (blue channel and violet laser) were used as display because the signal emitted by the secondary antibody coupled to Alexa Fluor™ Plus 405 was considerably higher than autofluorescence.

For secretion measurement, beads fluorescence was assessed by flow cytometry. Data were processed in Python with gating based on forward and side scatter to remove remaining cells (Supplementary Note 12). To account for variability in bead size, the ratiometric mNeonGreen/Cerulean signal was computed by dividing GRN-B values by BLU-V values. Then, the ratiometric signal given by Cerulean alone was subtracted to obtain the fluorescence coming from captured mNeonGreen.

### Confocal microscopy

For untreated samples, 6 µl of cultures are added in wells of an 18-well Ibidi µ-slide (CLINISCIENCES Ref. 81821) and mixed with 20 µL of PBS pH 7.5. For cells labelled with dye or antibodies, 26 µL of the final cell solution are transferred into a well of an 18-well Ibidi µ-slide. Images were acquired with BC43 spinning disc microscope (Oxford Instruments). All images were processed with Fiji macros using the same linear adjustment of pixel values for all samples of an experiment.

### Fluorescence Activated Cell Sorting

Culture samples from OptoTube experiments were resuspended in cold PBS pH 7.5 at OD600 0.75 (7.5×10^6^ cells/mL). Samples were transferred to a test tube with strainer snap cap and kept on ice before sorting. The sorting was performed with a BD FACSAria III (BD Biosciences) using the 561 nm laser. For each sample, 10000 events were acquired to define gating thresholds. FSC-A, SSC-A and FSC-H were used to gate cells (Supplementary Note 10). Then, the region containing 0.3% of the cells with the highest fluorescence detected through the 610/20 bandpass filter after excitation by the 561 nm laser was defined. Then, samples were subjected to sorting, and more than 10^6^ cells were assessed for each sample. Sorted and unsorted populations were grown overnight in liquid YPD at 30°C. Cultures were diluted several times and plated on solid YPD medium supplemented or not with hygromycin at 600 μg/mL. Colonies were counted using a counting pen (Thermo Scientific Ref. 11862710).

### Deep amplicon sequencing

First, 2.8 mL of cultures sampled from the eVOLVER bioreactors were used for genome extraction (Biosearch Technologies Ref. MPY80200). Then, a locus-specific PCR was done with the primer pair P998/P999, targeting the promoter and the 5’ of the scFv gene using the extracted genome as template. The PCR was performed with the high fidelity Phusion DNA polymerase (Thermo Scientific Ref. F530L). A 50 µL reaction was prepared with a primer concentration of 0.1 µM and 64 ng of genomic

DNA. The PCR program contained 22 cycles with an annealing temperature of 59.8°C and an elongation time of 1 minute 15 seconds. PCR products were purified with a PCR Clean-up kit (Macherey Nagel Ref. 740609.240C) using a Water/NT1 buffer ratio of 3/1. Second, the indexing PCR was performed following the Illumina protocol for 16S Metagenomic sequencing. The reaction was prepared in 25 µL total volume with the 2x Kapa Hifi HotStart ReadyMix (Roche Diagnostics Ref. 7958935001). 2.5 µL of the purified product of the locus-specific PCR were used as template. 5 µl of primers with Unique Dual Indexes were used (Illumina Ref. 20091660, set D). The PCR program contained 8 cycles with an annealing temperature of 55°C and an elongation time of 30 seconds. AMPure XP Beads for DNA Cleanup (Beckman Coulter Ref. A63882) were used for PCR product purification using 0.8X magnetic bead ratio. Purity of purified PCR products was assessed by running samples on the Fragment Analyzer using the High Sensitivity DNA/NGS Kit (Agilent Ref. DNF-474-0500).

Sequencing was performed on MiSeq i100 using the Series 5M Reagent kit 300 cycles (Illumina Ref. 20126565). Samples were pooled in equimolar quantities and mixed with 10% of PhiX (Illumina Ref. 15017872). Single-read sequencing was performed with 301 cycles, followed by 10 cycles for each index. Raw data were processed using the packages bowtie2 (indexing and read mapping) and samtools (filtering, sorting and counting).

### Digital PCR

Digital PCRs were performed with QIAcuity Probe PCR Kit (Qiagen Ref. 250102) in QIAcuity Nanoplate 26k 24-well (Qiagen Ref. 250001) with the QIAcuity One 5-plex device (Qiagen Ref. 911021). The experiment for the calibration curve was done in QIAcuity Nanoplate 8.5k 24-well (Qiagen Ref. 250011). Assays were done in duplex to quantify in the same sample the quantity of HygR gene (Custom dPCR Microbial assay, Qiagen Ref. 250208) and *COG6* gene (dPCR Microbial assay, Qiagen Ref. 250207, Id. DMA00758-H), following manufacturer instructions. Analyses were done on the software of the manufacturer, the same channel-specific thresholds were set for all samples using blank samples as negative control.

### Statistics and reproducibility

No statistical method was used to predetermine sample size. No data were excluded from the analyses. The experiments were not randomized. The Investigators were not blinded to allocation during experiments and outcome assessment.

## Data availability

Plasmid sequences used to construct all yeast strains are available in GenBank format in the AutoGrowth Git repository (https://gitlab.inria.fr/InBio/Public/AutoGrowth). Raw experimental data have been deposited on Zenodo (doi:10.5281/zenodo.21282497), they include: flow cytometry data (cell fluorescence, secretion, and display), microscopy images, OD data from plate reader, digital PCR data, deep sequencing data, colony counts, bioreactors data (pump activation, automated flow cytometry data, and OD data).

## Code Availability

All scripts have been deposited in the AutoGrowth git repository (https://gitlab.inria.fr/InBio/Public/AutoGrowth). We provide (i) notebooks ran for experiments performed with the ReactSight bioreactor platform, (ii) notebooks and scripts used to process raw data and generate figures, (iii) Fiji macros for microscopy image analysis.

## Acknowledgements

The authors would like to thank Tom Ellis for sharing the yWS677 strain and providing sound advice, along with Pierre Laffaye for the sequence of the anti-NS1 nanobody 21 (Nb21). We are grateful to Sean Kennedy, François Bertaux and Sebastián Sosa-Carillo for their valuable input. We thank the Single Cell Biomarkers and the Photonic BioImaging units of technology and service at the Institut Pasteur for support in conducting this study. We also acknowledge the support of the Biomics (Laurence Ma) and Flow Cytometry (Sébastien Megharba) Platforms at Institut Pasteur.

This work was supported by ANR grants SmartSec (ANR-21-CE44–0033) and TrojanYeast (ANR-24-CE18–2885), by Region Île-de-France in the framework of DIM BioConvS (PlatPath grant), and by the Inria/IFPEN joint laboratory (Screen2Learn grant).

## Author contributions

H.G. and G.B. conceived the study. H.G. constructed all strains and performed most of the experimental work. C.C. and S.N. developed the eVOLVER-based ReacSight platform. A.F.D.S. developed the yeast surface display protocol. H.G. performed the data analysis. G.B. and S.N. supervised the study. H.G. and G.B. wrote the manuscript with input from S.N.. S.N. and G.B. contributed equally to the work.

