## Supplementary Notes for "Coupling cell growth to protein secretion enables pooled screening at single-cell resolution in bioreactors"

### Supplementary Note 1: Strains, plasmids and proteins of interest used in this study

#### Strains and plasmids

Double click on the icon to open associated file: AutoGrowth_strains.xlsx

Of note, most Level 0 plasmids came from plasmid kits: Yeast GPCR-sensor Toolkit (Addgene Ref. 1000000157), Multiplex Yeast Toolkit (Addgene Ref. 1000000229), and EasyClone-MarkerFree Vector Set (Addgene Ref. 1000000098). Sequences of the others are provided in the file.

#### Properties of the POIs tested

***Table S1**: Proteins of interest and their properties.*

| **Short name** | **POI under study** | **Native organism** | **GenBank ID** | **PTM*** | **Size (aa)**** |
|---|---|---|---|---|---|
| mNG | mNeonGreen[1] | n.a. | BBB44438 | n.a. | 382 |
| XylC | Endo-1,4-beta-xylanase C[2] | *Aspergillus niger* | EU848304 | 1x S-S | 690 |
| Xyl2 | Endo-1,4-beta-xylanase 2[3] | *Trichoderma reesei* | P36217.2 | 2x N-glyc | 572 |
| Nb21 | Anti-NS1 nanobody 21[4] | n.a. | n.a. | 1x S-S | 527 |
| scFv | Single chain variable fragment 4M5.3[5] | n.a. | 1X9Q_A | 2x S-S | 637 |
| Amy | α-amylase[6] | *Aspergillus oryzae* | CAA31220 | 4x S-S + 2x N-glyc | 860 |

* **PTM**: post-translational modifications; **S-S**: disulfide bond; **N-glyc**: N-glycosylation.

** Includes the protein of interest, the MFo2 signal peptide, the fluorescent reporter, the tag (3xFLAG 1xHis) and the α-factor.

### Supplementary Note 2: Assessing the proper functioning of the optogenetic induction system for displayed and secreted mNeonGreen proteins

Green fluorescent signal increases with induction for both secreted and displayed mNeonGreen. Two phenotypes are present for secreted mNeonGreen at 66% and 100% light duty cycles, with cells showing fluorescence in the vacuole or in the cytoplasm. For displayed mNeonGreen, there is a clear GFP signal at the cell surface, which increases with induction, while there is also a weaker signal from the vacuole. In agreement with the flow cytometry data (Figure 1C), there is no blue fluorescent signal (*i.e.* display signal) for secreted mNeonGreen while there is clear blue signal for displayed mNeonGreen which increases with induction (*i.e.* light duty cycles).

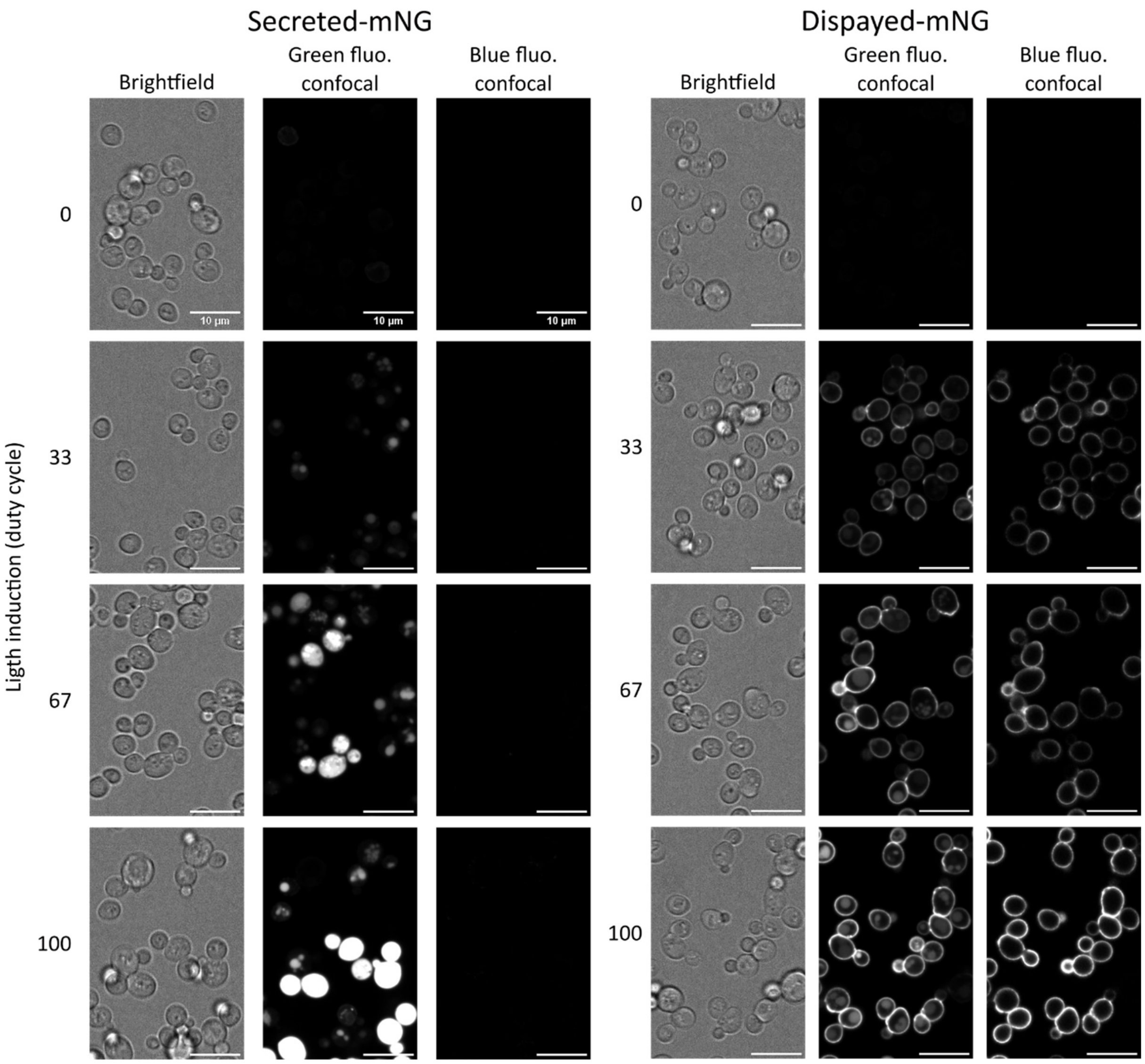

***Figure S2.1***: *Confocal microscopy images of cells expressing secreted or displayed mNeonGreen after 10h of culture under various duty cycles of blue light illumination with the OptoTube setup, and antibody labeling. Green and blue fluorescence are emitted by mNeonGreen and the secondary antibody used for display measurement, respectively.*

Interestingly, expressing secreted mNeonGreen at 100% duty cycle altered cell fitness, as the OD reached at the end of the culture was lower. This was not observed for displayed mNeonGreen.

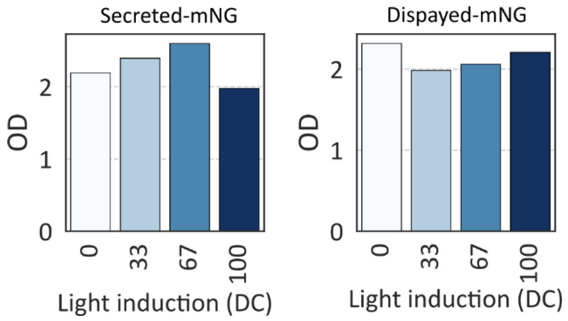

***Figure S2.2**: OD measurements of cells expressing secreted or displayed mNeonGreen after 10h of culture under various duty cycles of blue light illumination with the OptoTube setup.*

### Supplementary Note 3: Proving the cell surface activation of Ste2id and the importance of α-factor fusion

We observed that secreted mNeonGreen causes strict autocrine activation, that is, triggers no paracrine activation (Figure 2A). GPCRs, such as Ste2, can in principle be activated inside the cell. This could explain the observed strict autocrine activation. This would not be a desired feature of our sensing system. Indeed, downstream activation should reflect the quantity of secreted proteins. So, to investigate this, we assessed the localization of the activation. Having validated that the activation of the Ste2id receptor takes place mostly at the cell surface, we investigated the surprising absence of paracrine activation for secreted mNeonGreen fused to α-factor. To do so, we tested paracrine signaling using three different "POIs": α-factor alone, tag-α-factor, and mNeonGreen-tag-α-factor.

#### Cell surface activation of Ste2id

dTA, an antagonist of α-factor, inhibits activation of Ste2 at the cell surface, thereby allowing us to interrogate whether POI-based downstream activation takes place at the surface, as expected. We first demonstrated that dTA inhibits the activation of Ste2id variant by α-factor (Figure 3.1A, left). Then, we found that POI-based downstream activation is also reduced upon dTA addition (Figure S3.1A, right), suggesting that Ste2id binding does take place mainly at the cell surface. However, there seems to be residual activation. A KREAEA protection, cleaved by Kex2 protease in the trans Golgi network (TGN), and a LLEAEA protection, uncleavable, were fused at the C-terminal of α-factor to localize residual activation (Figure S3.1B). The lack of activation seen with the uncleavable protection indicates that cleavage is necessary for Ste2id activation. mNeonGreen harboring the cleavable protection mimics unprotected POI activation. This means that activation of Ste2id takes place after the Kex2 cleavage, in the late stages of secretion, either in the TGN, in secretory vesicles, or outside of the cell.

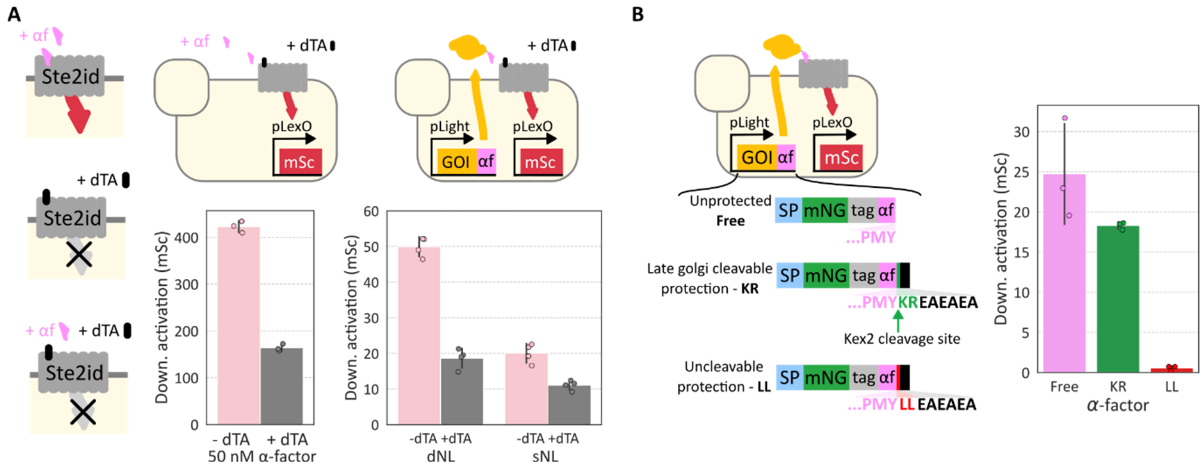

***Figure S3.1**: Study of the localization of Ste2id activation. **A)** Effect of dTA on downstream activation caused by either 50 nM α-factor (left), or secreted/displayed mNG-α-factor induced with 100% blue light duty cycle in the OptoTube setup (right). **B)** Autocrine strains secreting mNG with or without an α-factor protection were induced with 100% light duty cycle for 10 hours in the OptoTube setup before measurement of cell fluorescence by flow cytometry. Experiments are done in biological triplicate; medians of each population are represented by dots, and their mean and the corresponding standard variation is*

**Importance of α-factor fusion to prevent paracrine activation**

As most of the activation takes place at the cell surface, we would expect secreted mNeonGreen released in the medium to activate other cells (*i.e.* paracrine activation). We assessed such autocrine and paracrine activation using co-cultures. As in Figure 2A, they were composed of an autocrine strain and a paracrine strain. However, here, autocrine strains secrete different "POIs": α-factor (sAL), tag-α-factor (sTL), and mNeonGreen-tag-α-factor (sNL). We used only "Low sensitivity" autocrine strains because their operational range is larger, which avoids saturation of the response. As expected, secreted α-factor produces a very strong autocrine activation. Surprisingly, activation given by sTL is similar to the one given by sNL. This holds true irrespective of the type of paracrine cells used (Low or High sensitivity strains, cBL or cBH). This means that fusing only a Flag-His tag at the N-terminal of α-factor significantly alters its ability to generate paracrine activation. This agrees with a previous study showing that mutations in the N-terminus of α-factor could affect paracrine activation, but not autocrine activation[7].

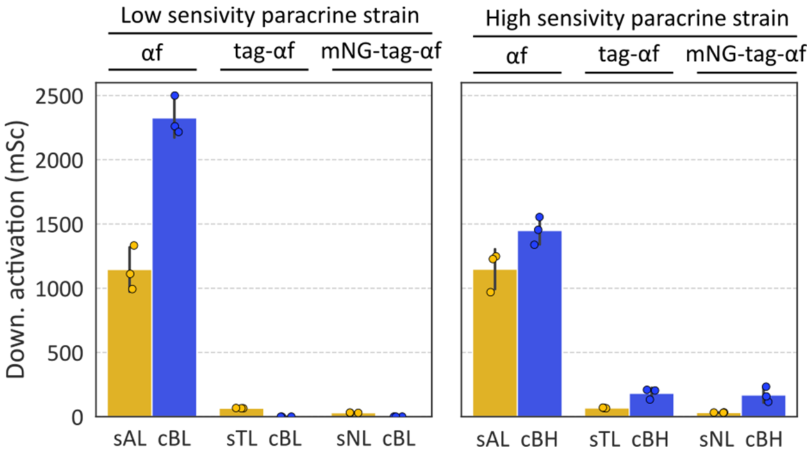

***Figure S3.2**: Co-culture of various autocrine and paracrine strains grown under 100% light duty cycle for 10 hours, before measurement by flow cytometry of cell fluorescence. Experiments are done in biological triplicate; medians of each population are represented by dots, and their mean and the corresponding standard variation is shown with the bar plot.*

### Supplementary Note 4: Growth rate computation in batch and continuous cultures

#### Batch cultures

The determination of growth rates in batch cultures was performed in the same way for OptoPlate and OptoTube experiments (Figure S4.1). The growth rate $\mu$ of cells in exponential phase can be inferred with a linear regression on the logarithm of the density of cells $X_t$:

$$(1.1) \qquad X_t = X_0 e^{-\mu t} \quad \leftrightarrow \quad \log(X_t) = \mu t + \log(X_0)$$

Three OD timepoints were taken, and their logarithmic values fitted to linear function using least-squares regression. We did simulations using the inferred growth rates and observed that data points and simulations are matching. Besides confirming the relevance of the inferred growth rate values, it confirms that cells were growing exponentially.

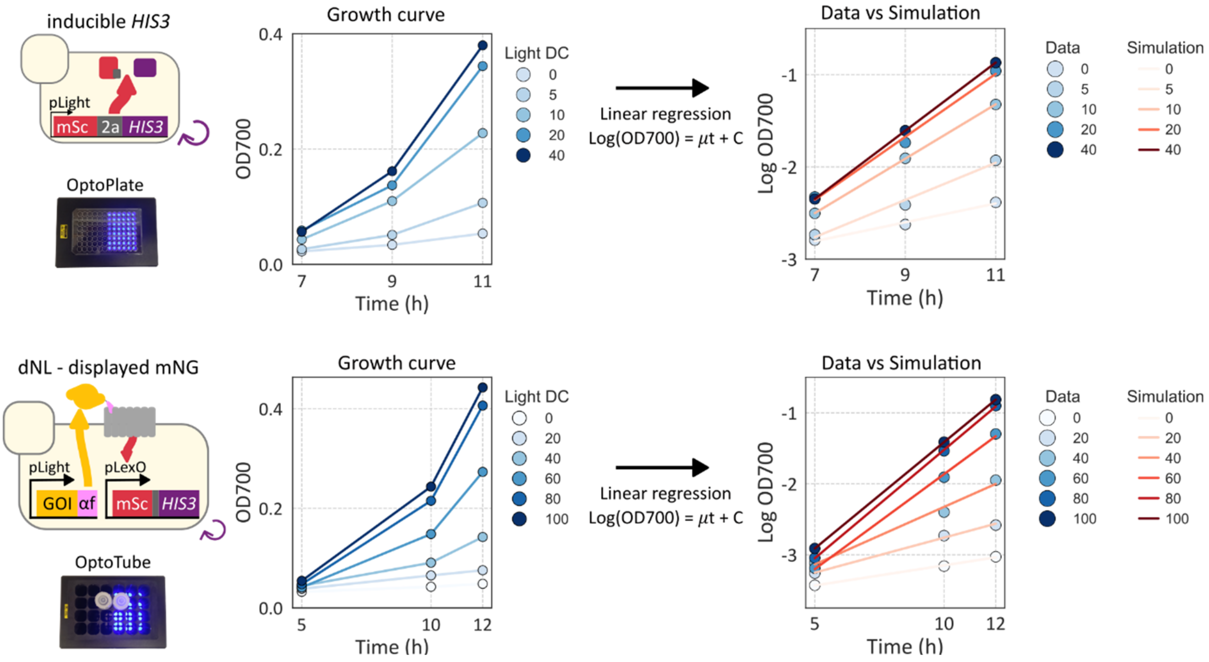

***Figure S4.1***: *Cells grown in OptoPlate and OptoTube under various light duty cycles are sampled at three different time points and their OD measured using a plate reader. A linear regression is performed on the logarithm of the OD value to infer the growth rate.*

**Continuous culture**

We conducted continuous culture experiments under the turbidostat regime, in which the OD is kept constant at 0.5. Keeping a constant OD results in equivalence between the growth rate $\mu$ and the dilution rate $D$:

$$(2.1) \quad \frac{dOD}{dt} = \mu \times OD - D \times OD$$

$$(2.2) \quad 0 = \mu \times OD - D \times OD$$

$$(2.3) \quad \mu = D$$

The dilution rate $D$ represents the relationship between the volume of the culture $V$ and the flow of medium into the culture $F$. Hence, growth rate $\mu$ can be expressed in function of $F$ and $V$:

$$(3.1) \quad D = \frac{F}{V} \rightarrow \mu = \frac{F}{V}$$

In our case, the volume of culture $V$ is constant, so the growth rate $\mu$ is proportional to the flow of medium $F$. To avoid relying on pump calibration, we use pump activation count as proxy of the medium flow. It is possible because a given pump adds the same volume at each activation. This makes the number of pump activations over a given time window proportional to the added volume. We use a time windows of 4 hours to obtain smooth signals that can be interpreted. In all bioreactor experiments, cells are grown in non-selective medium in the dark for several hours before the start of induction. We use these pre-induction windows to normalize the activation count, so that bioreactors can be compared between them.

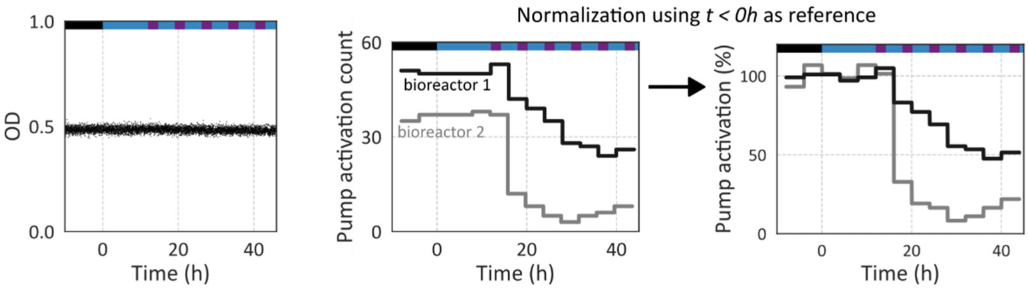

***Figure S4.2**: Computation of pump activation as a proxy for growth rate in continuous culture leveraging the fact that constant OD results in growth rates being equivalent to dilution rates.*

### Supplementary Note 5: Cell surface accumulation pattern depends on the nature of the secreted protein

For each POI, we showed that the relationship between secretion and downstream activation is monotonic, except in cases where strong POI induction alters cell fitness. We represented the same data on a single plot and used logarithmic scale for more clarity (Figure S5.1). Taking one POI at a time, we observe again the monotonic relationship between secretion and downstream activation, except for Nb21. However, when comparing POIs between them, we observe that highly secreted proteins have lower downstream activation. This is rather counterintuitive. We hypothesized that efficiently secreted POIs transit rapidly through the secretory pathway and cell wall, whereas poorly secreted POIs accumulate along the late stages of the secretory pathway, in the periplasm or in the cell wall, and are thus more likely to activate Ste2id. In other words, high flux through the secretion pathway would lead to low local concentrations of secreted proteins, while low flux would lead to high local concentrations.

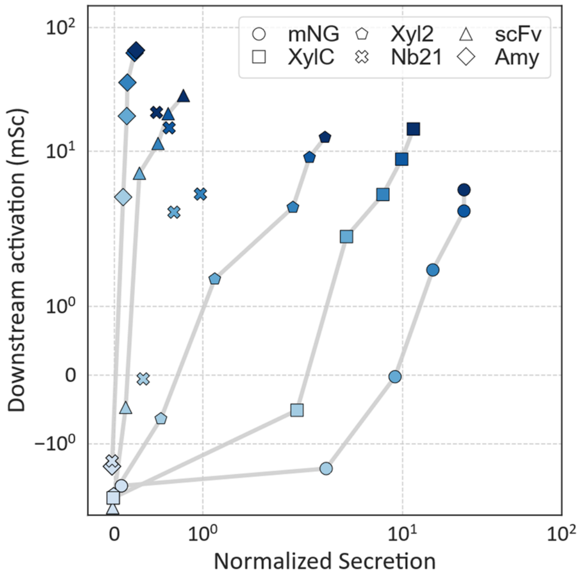

***Figure S5.1***: *Hard-to-secrete POIs show higher downstream activation levels. Autocrine strains secreting various POIs are grown in his+ medium under a range of blue light for 12h, before inoculation of fresh his+ medium for 10 hours, followed by flow cytometry measurement of cell fluorescence and secretion, through anti-Flag magnetic beads.*

We took advantage of the mNeonGreen tag to investigate the subcellular localization of the POIs (Figure S5.2). Some of these images are present in the main text, but we added here images of cells secreting Xyl2 and Nb21. Xyl2 accumulates in the vacuole, as the other POIs, while Nb21 shows strong cytoplasmic accumulation, a possible reason for its deleterious effect on cell fitness. These phenotypes are indicative of overwhelmed secretory pathways. Vacuolar accumulation comes from protein mis-sorting in the TGN or ER-phagy, while cytoplasmic accumulation is most likely originating from saturation of the ER import machinery. As discussed in the main text, signal at the cell periphery is observed for scFv and Amy only. This signal is present, and for some cells stronger, at the bud neck, confirming that it comes from the periplasm and not from the cortical ER. Therefore, scFv and Amy accumulate at the cell surface.

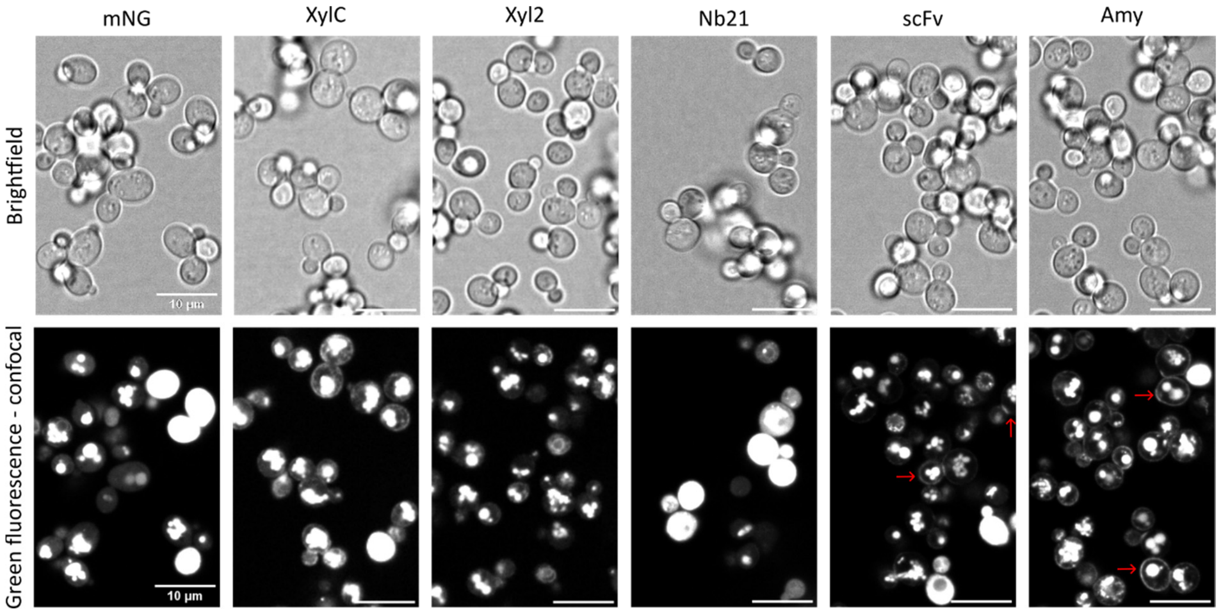

***Figure S5.2**: Signal at the cell periphery observed for scFv and Amy only. Autocrine strains are grown in his+ medium under 100% blue light DC for 12 hours, before inoculation of fresh his+ medium for 10 hours and transfer of samples into wells of ibidi µ-slide for confocal microscopy (60x).*

### Supplementary Note 6: Capacity of the biosensor to generate scFv secretion-dependent growth in bioreactors

We decided to use scFv for proof-of-concept experiments, and started by characterizing the strain in bioreactors. We wanted to confirm in bioreactors the results obtained in batch about the correlation between downstream activation and scFv expression level. We also assessed whether increased downstream activation translates into higher growth rate in selective medium. To investigate these questions, we induced cells with a range of light duty cycles and grew them in 40% histidine medium before medium switch to his- medium at 12h and his- 50 µM 3AT at 24h (Figure S6.1). As expected, graded scFv induction leads to graded downstream activation, confirming results obtained in batch. In addition, we observed that the switch to his- medium generates graded growth rates, and that, increasing stringency by 3-AT addition, further reduces growth rates. We investigated the relationship between downstream activation and growth rate in both media. In his- medium, growth rate scales with downstream activation for induction between 0 and 60%, a threshold after which growth rate seems to plateau at around 75% of pump activation. The fact that a plateau is reached below 100% pump activation probably comes from the burden imposed on the cell fitness by strong secretion. The increased stringency caused by 3AT addition results in a monotonic increase of growth with downstream activation for the full induction range. This graded relationship suggests that the biosensor should support screening for improved scFv secretion.

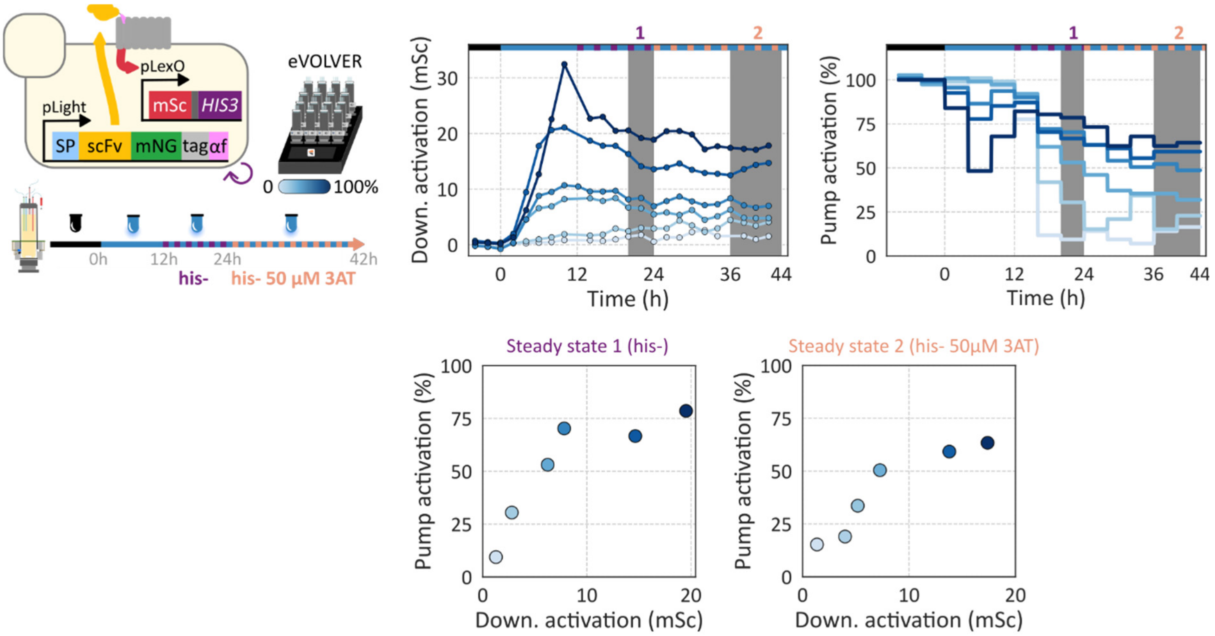

*Figure S6.1: The biosensor permits the growth rate to scale with scFv induction in selective medium. Autocrine strain secreting scFv was grown in turbidostat regime at OD 0.5 in 40% histidine medium in dark, to allow cells to reach exponential growth. Then, various blue light duty cycles were applied in his+ medium for 12 hours before switching to his- medium, and later switch to his- 50 µM 3AT at 24h post light induction. Cell fluorescence is measured every 2 hours by automated flow cytometry and median fluorescence of each population is shown here. Steady state values are obtained by taking the mean from either the window 20-24h or the last 8 hours of culture.*

Of note, the transient decay of growth rate observed between 0h and 12h for strong induction was reported previously[8]. In brief, strong scFv induction results in a bimodal population, with some cells showing high intracellular accumulation of the protein. This transient bimodality is associated with the growth rate decay observed at the population level. The intracellular accumulation of the transient population most likely corresponds to the cytoplasmic accumulation observed in Figure S5.2. Although this growth rate decay is transient, the pre-induction growth rate is not totally recovered after. As discussed above, it probably comes from the burden associated with strong scFv induction.

### Supplementary Note 7: Impact of signal peptides on the trafficking of secreted scFv

***Table S7**: Signal peptides tested for scFv secretion.*

| Short name | Full name | Gene | References |
|---|---|---|---|
| MF8 | MFαprepro8 | *MF(alpha)1* | [9,10] |
| OsMF2* | OST1pre-MFαpro | *OST1 / MF(alpha)1* | [11,12] |
| MFo2 | MFαpreproOPT | *MF(alpha)1* | [13] |
| OsMF | OST1pre-MFαpro | *OST1 / MF(alpha)1* | [10,14] |
| FLO10* | | | [15] |
| SWP1* | | | [15] |
| SUC2* | | | [12] |
| FET3* | | | [15] |
| MID2* | | | [15] |
| MF* | MFαprepro | *MF(alpha)1* | [12] |
| YT* | Yap3-TA57 | *YAP3* | [12,16] |
| GAS5* | | | [15] |
| MFo | MFαpreproOPT | *MF(alpha)1* | [10,17] |

* codon remastered, still using codon-usage of *S. cerevisiae*, and extra Methionine in N-terminal

We compared the amino acid sequences of the SP based on the signal peptide of the *MF(alpha)1* from the S288C strain, called MF_S288c. The DNA sequences of MF and MF_S288c are different but they encode the same protein sequence. By contrast, the optimized versions MFo, MFo2, and MF8 contain mutations at the protein level:

- MFo has mutations on the residues A9D, A20T, L42S, and D83E,
- MFo2 is different at L42S, and with the C-terminal motif KREEGEPK replacing the KREAEA motif,
- MF8 has mutations on residues V22A, G40D, L42S, V50A, V52A, F55L, L64S, F65S, I66T, S81Q.

```
MF8           MRFPSIFTAVLFAASSALAAPANTTTEDETAQIPAEAVIDYSDLEGDFDAAALPLSNSTN
MFo           MRFPSIFTDVLFAASSALATPVNTTTEDETAQIPAEAVIGYSDLEGDFDVAVLPFSNSTN
MF_S288c      MRFPSIFTAVLFAASSALAAPVNTTTEDETAQIPAEAVIGYLDLEGDFDVAVLPFSNSTN
MF            MRFPSIFTAVLFAASSALAAPVNTTTEDETAQIPAEAVIGYLDLEGDFDVAVLPFSNSTN
MFo2          MRFPSIFTAVLFAASSALAAPVNTTTEDETAQIPAEAVIGYSDLEGDFDVAVLPFSNSTN
              ******** **********:*.*****************.* *******.*.**:*****

MF8           NGLSSTNTTIASIAAKEEGVQLDKREAE---
MFo           NGLLFINTTIASIAAKEEGVSLEKREAE---
MF_S288c      NGLLFINTTIASIAAKEEGVSLDKREAEA--
MF            NGLLFINTTIASIAAKEEGVSLDKREAEA--
MFo2          NGLLFINTTIASIAAKEEGVSLDKREEGEPK
              ***   **************.*:***
```

***Figure S7.1**: Alignment of the SP evolved/mutated from MF(alpha)1 and the endogenous one (MF_S288c). It was done with the Multiple Sequence Alignment tool from Clustal Omega (EMBL-EBI).*

We also provide a comparison of OsMF and OsMF2. Of note, OsMF2 bears a point mutation (G244T) at the DNA level, causing the missense mutation G82C at the protein level compared to the sequence from Durmusoglu *et al.*[12]. They otherwise have the same amino acid sequence, except for the positions 45, 86, and 92. However, their encoding varies with silent mutations in 32 codons out of 92.

```
OsMF     MRQVWFSWIVGLFLCFFNVSSAAPVNTTTEDETAQIPAEAVIGYLDLEGDFDVAVLPFSN
OsMF2    MRQVWFSWIVGLFLCFFNVSSAAPVNTTTEDETAQIPAEAVIGYSDLEGDFDVAVLPFSN
         ******************************************** ***************

OsMF     STNNGLLFINTTIASIAAKEEGVSLDKREAE-
OsMF2    STNNGLLFINTTIASIAAKEECVSLEKREAEA
         ********************* ***:*****
```

***Figure S7.2****: Alignment of the SP developed from the pre-pro region of MF(alpha)1 and OST1 genes. It was done with the Multiple Sequence Alignment tool from Clustal Omega (EMBL-EBI).*

We assessed individually in OptoTube the 13 autocrine strains, each secreting scFv with a different SP. As described in the main text, downstream activation is scaling with secretion. On the contrary, we observe in Figure S7.3 that internal POI, meaning intracellular scFv, is higher for inefficient SPs, with the exception of MF8, promoting accumulation despite good secretion level. Interestingly, the 6 SPs taken from Xue et al.[15] have the same rank of secretion efficiency here as in their study on α-amylase secretion, with the exception of GAS5, which was the best in their case.

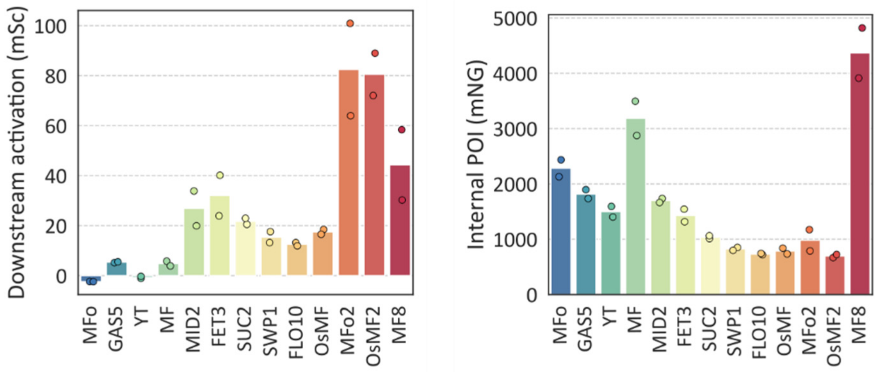

***Figure S7.3****: Efficient signal peptides tend to generate lower intracellular accumulation, except MF8. Autocrine strains secreting scFv with various SPs are grown in OptoTube setup in his+ medium for 4 hours in the dark, then for 12 hours under 100% blue light DC, before inoculation of fresh his+ medium for 10 hours under blue light followed by flow cytometry measurement of cell fluorescence. Experiments are done in biological duplicates; medians are represented with dots, and their means shown by the bar plot.*

We investigated further the differences caused by SPs on scFv trafficking through the secretory pathway by looking at the subcellular localization with confocal microscopy after growth under maximal induction (100% light duty cycle) in the OptoTube setup. We first observed that the subcellular localization of secreted scFv was very dependent on the SP. Interestingly, inefficient SPs (10 to 13) tended to cause accumulation in both the cytoplasm and in the nuclear ER. Intermediate SPs (4 to 9) seemed to cause accumulation in both nuclear and cortical ER, the latter being hard to distinguish from a potential cell surface signal. Except MF8, efficient SPs (1 to 3) caused scFv to accumulate in the vacuole and at the cell surface, and a weak nuclear ER signal was present in few cells only. Microscopy results confirmed flow cytometry data with higher accumulation of scFv in cells/constructs with inefficient SPs and MF8.

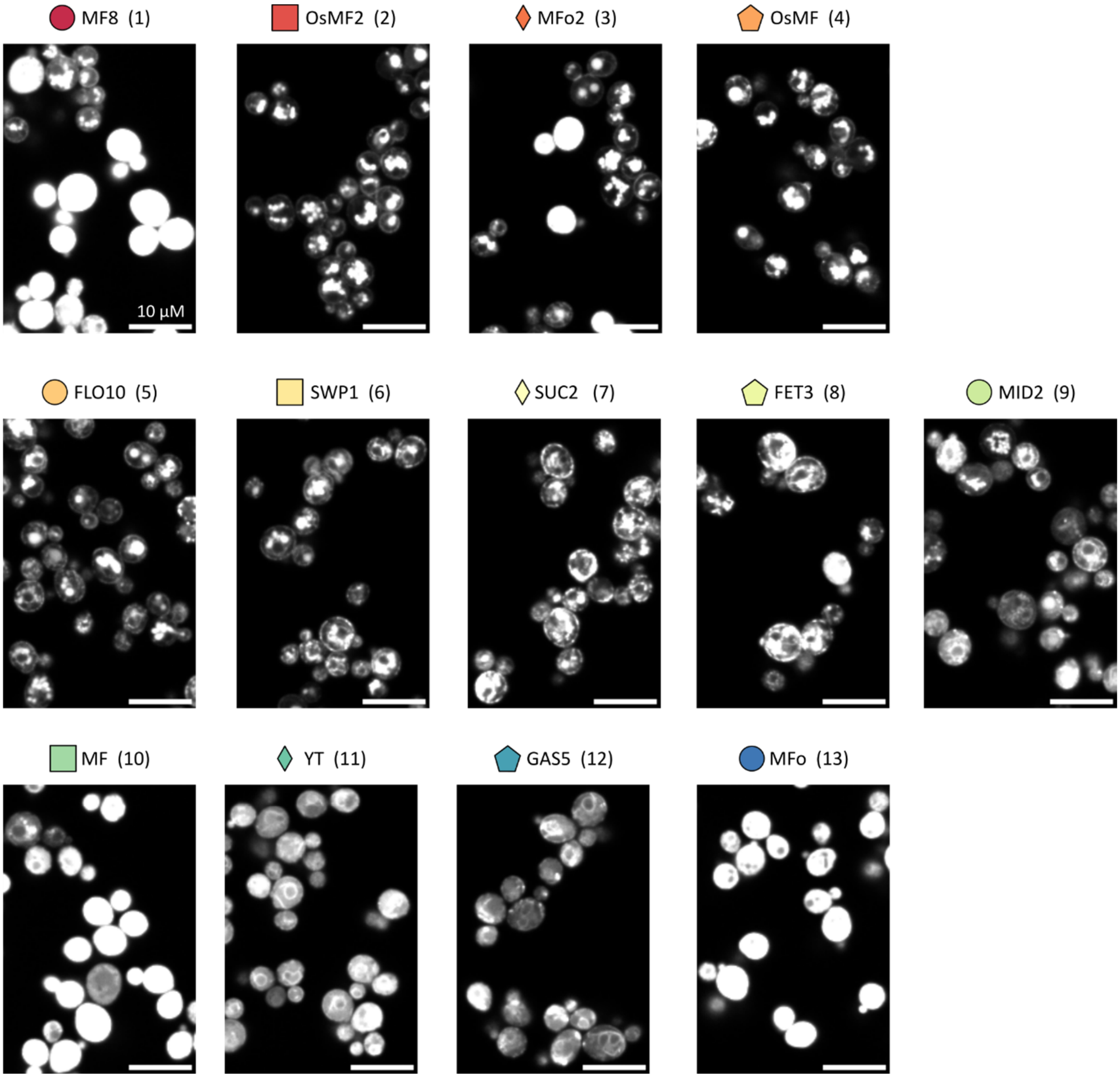

***Figure S7.4**: Impact of SP on scFv trafficking through the secretory pathway. Autocrine strains secreting scFv with various SP are grown in his+ medium under 100% blue light DC for 12 hours, before inoculation of fresh his+ medium for 10 hours and transfer of samples into wells of ibidi µ-slide for confocal microscopy (60x).*

### Supplementary Note 8: Capacity of the biosensor to identify the effect of genetic perturbations on the scFv secretion

The optimization of secretion can be achieved by engineering the endogenous gene network. We set out to test whether the biosensor would be able to identify the effect of genetic perturbations, deletion or overexpression, on scFv secretion. We selected 8 genes whose deletion or overexpression were shown previously to increase secretion of recombinant proteins.

***Table S8**: Genes whose deletion or overexpression were tested on the scFv secretion.*

| Gene | Function (from https://www.yeastgenome.org/) | Modality | Ref. |
|---|---|---|---|
| *HDA2* | Subunit of HDA1 histone deacetylase complex. | deletion | [18] |
| *SEC65* | Subunit of the signal recognition particle (SRP); involved in protein targeting to the ER. | overexpression | [19] |
| *PDI1* | Protein disulfide isomerase; multifunctional oxidoreductase of the ER lumen, essential for disulfide bond formation in secretory and cell-surface proteins, and processing non-native disulfide bonds. | overexpression | [20–24] |
| *KAR2* | ATPase involved in protein import into the ER; also acts as a chaperone to mediate protein folding in the ER and may play a role in ER export of soluble proteins. | overexpression | [20,22,23] |
| *CWH41* | Processing alpha glucosidase I; ER type II integral membrane N-glycoprotein involved in assembly of cell wall beta 1,6 glucan and asparagine-linked protein glycosylation; has a role in ER protein quality control and sensing of ER stress. | overexpression | [24,25] |
| *SSO2* | Plasma membrane t-SNARE; involved in fusion of secretory vesicles at the plasma membrane. | overexpression | [26] |
| *SNC2* | Vesicle membrane receptor protein (v-SNARE); involved in the fusion between Golgi-derived secretory vesicles with the plasma membrane. | deletion | [18] |
| *VPS5* | Nexin-1 homolog; required for localizing membrane proteins from a prevacuolar/late endosomal compartment back to late Golgi. | deletion | [27] |

For deletions, the entire ORF was removed with CRISPR/Cas9-mediated editing. For overexpression, we placed the genes under the doxycycline (dox) inductive pTET promoter. In both cases, the secretion of scFv was assessed under 100% blue light DC in the OptoTube setup. We observed a range of secretion levels, with most perturbations increasing secretion compared to the parental strain. Previously seen beneficial, the deleterious effect of *CWH41ox* and *hdaΔ* could come from the dependency of these perturbations on the experimental conditions. s*nc2Δ* grew poorly, with a final OD approximately twice lower than others. Downstream activation scaled with secretion, with the exception of *PDI1ox*, showing low activation despite high secretion. It could be an example of secretion improvement through better diffusion through the cell wall as Pdi1 facilitates proper folding of the POI. Noteworthy, *vps5Δ* exhibits 4- and 6-fold increases in secretion and downstream activation, respectively.

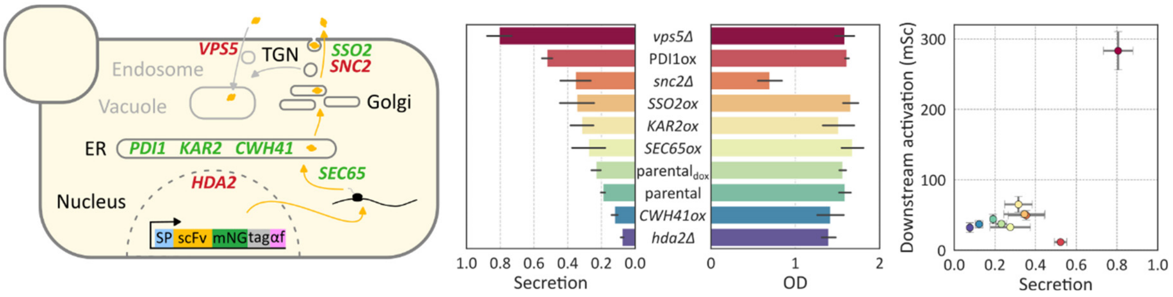

***Figure S8.1***: *The biosensor can be used to identify beneficial genetic perturbations. Autocrine strains secreting scFv, and deleted for a gene, are grown in his+ medium for 4 hours in the dark, then for 12 hours under 100% blue light DC, before inoculation of fresh his+ medium for 10 hours under blue light, followed by flow cytometry measurement of cell fluorescence and secretion, through anti-Flag magnetic beads. Strains with overexpression of endogenous genes were subjected to the same protocol, with the exception that medium was supplemented with 1 µg/mL of doxycycline (rtTA/pTET system used for overexpression). The parental strain was assessed with and without doxycycline. Experiments are done in 5 biological replicates; their mean and the corresponding standard variation is shown with the bar plots, and the scatter plot.*

### Supplementary Note 9: Effect of *VPS5* deletion in BY4741 background and its characterization in bioreactor

*VPS5* deletion was shown in Supplementary Note 8 to increase secretion of scFv. We sought to test whether this effect would translate in a BY4741 background, without an engineered mating pathway. Next, we compared downstream activation and growth rate of the parental and *vps5Δ* autocrine strains in bioreactors in selective medium. These experiments helped us to prepare and interpret the spike-in experiments of Figure 4B, notably by using a simple model of the co-culture.

**Effect of *VPS5* deletion on scFv secretion in BY4741 background**

Strains were assessed in the usual OptoTube setup with the same protocol as in Figure S8.1. Using the canonical BY4741 background, the deletion of *VPS5* caused a 3-fold increase of secretion, which is rather similar to the 4-fold obtained in the autocrine strain. It shows that the beneficial effect of the deletion transfers from the engineered background to the BY4741 background.

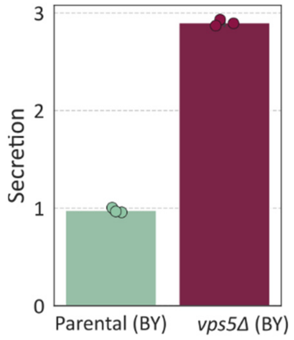

***Figure S9.1**: Parental and vps5Δ strains secreting scFv are grown in his+ medium for 4 hours in the dark then for 12 hours under 100% blue light DC using the OptoTube setup, before inoculation of fresh his+ medium for 10 hours under blue light followed by flow cytometry measurement secretion, through anti-Flag magnetic beads. Experiments are done in biological triplicate; individual samples are represented by dots, and their means are shown with the bar plot.*

**In-depth study of the *vps5Δ* autocrine strain**

We investigated whether the very high downstream activation of the *vps5Δ* strain would translate into faster growth in selective medium (Figure S9.2). Parental and *vps5Δ* strains secreting scFv were cultured in bioreactors with light induction at 0h and medium switch at 12h (his-) and at 24h (his- 0.1 mM 3AT). As seen in batch, *vps5Δ* strain showed much higher autocrine activation than the parental strain. This bioreactor experiment confirmed the stability of the signal. Growth of the two strains was similar until 16h, time at which the parental started to grow slower. Addition of 3AT at 24h further decreased the growth rate of the parental strain, while the growth rate of the *vps5Δ* strain remained unchanged. At steady state, growth rates of the parental and *vps5Δ* strains were 62% and 80% of the pre-induction growth rate, respectively.

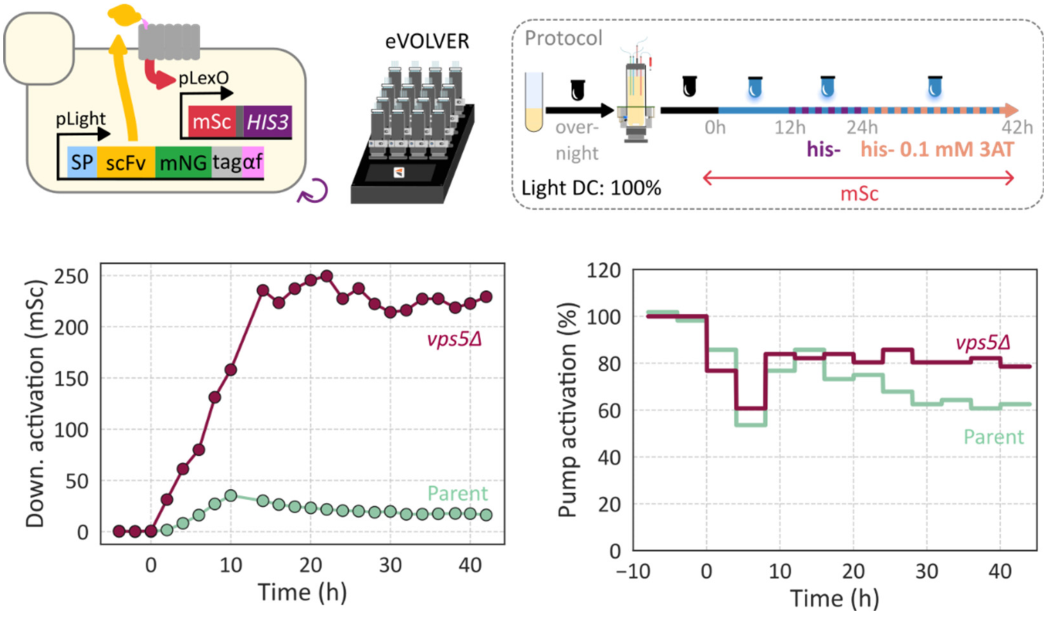

***Figure S9.2**: Autocrine parental and vps5Δ strains secreting scFv were grown in turbidostat regime at OD 0.5 in 40% histidine medium in dark to allow cells to reach exponential growth. Then, 100% blue light duty cycle was applied for 12 hours before switching to his- medium, and later switch to his- 0.1 mM 3AT at 24h post light induction. Cell fluorescence is measured every 2 hours by automated flow cytometry and median fluorescence of each population is shown.*

Next, we sought to simulate the enrichment of the *vps5Δ* strain in a co-culture with the parental strain in selective medium (his-, 0.1 mM 3AT), where the *vps5Δ* strain represents 0.1% at the culture onset. We used the steady state values, 62% and 80%, obtained previously, which are based on the pre-induction growth rate. We computed this pre-induction growth rate for each strain using OD data from the bioreactor experiments presented in Figure S9.2. Bioreactors were inoculated at OD 0.2 so that there was first a growth phase without dilution until the culture reached OD 0.5, value at which the turbidostat regime started (Figure S9.3A). We used this initialization phase to compute the growth by doing a linear fit on the log of the OD values (avoiding the post-inoculation lag phase). We found a growth rate of 0.36 $h^{-1}$ for both strains. We applied 62% and 80% to this value and obtained the predicted growth rate of both strains at steady state in selective environment (his-, 0.1 mM 3AT). Beginning with 0.1% of *vps5Δ* strain, we simulated the composition of the co-culture over 6 days, with 14 hours as the starting point, because this is the time of the switch to selective medium (Figure S9.3B). After 4 days of selection, at 110h, the *vps5Δ* strain was predicted to represent approximately 45% of the population, and above 90% after 6 days, which means respectively over 400- and 900-fold enrichments. Of note, the model assumes immediate change of the growth rate upon medium switch at 14 hours, whereas in reality the selective medium will take few hours to replace the 40% histidine medium. Hence, the growth rate change will take few hours so we can expect real enrichment dynamics to be slightly slower than in the simulation.

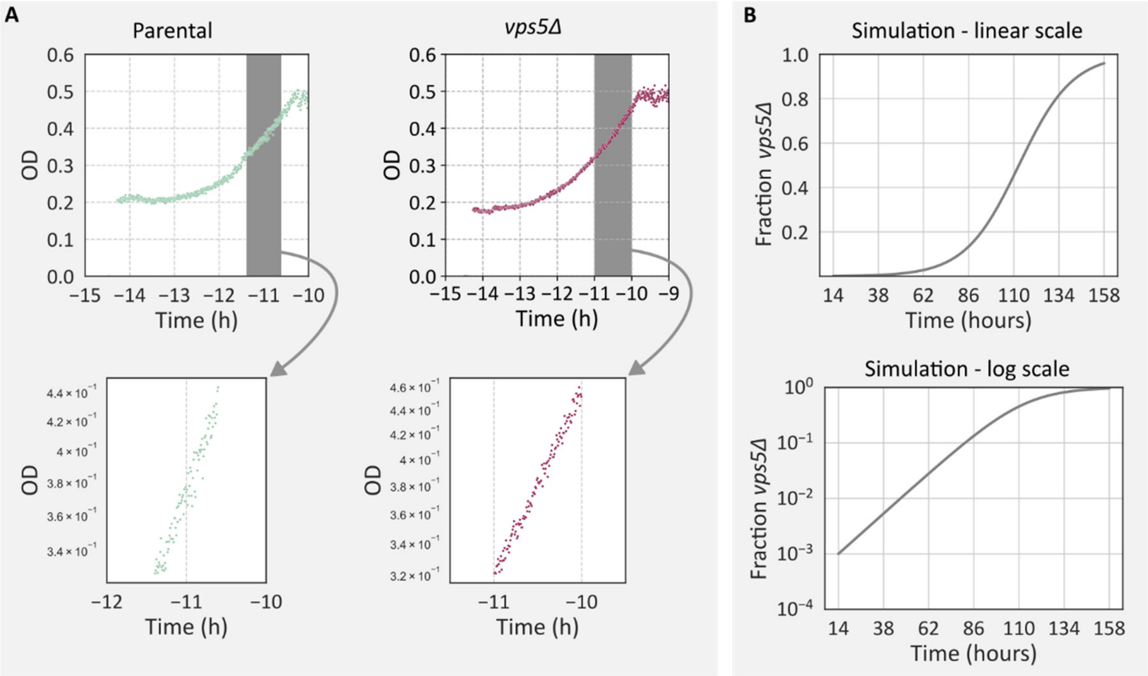

***Figure S9.3**: **A)** Autocrine parental and vps5Δ strains secreting scFv were grown in bioreactors, a 1-hour window before reaching OD 0.5 is used to compute the growth rate of the strains. **B)** Simulation of the vps5Δ cell fraction in a co-culture, starting with a vps5Δ:parental ratio of 1:1000, in continuous culture.*

### Supplementary Note 10: Computation of the enrichment of the *vps5Δ* strain in spike-in experiments

Spike-in experiments, presented in Figure 5B, were conducted to study the capacity of the biosensor to enrich the *vps5Δ* strain. This required computing the fraction of *vps5Δ* cells in the population. We did it using two procedures, plating and digital PCR, which we present in this note. We also provide enrichment results obtained using FACS.

**Plating procedure to compute fraction of *vps5Δ* cells**

For each sample of the spike-in experiments, we performed several 10-fold dilutions of the culture and plated 50 µL on YPD medium, supplemented or not with hygromycin. Several dilutions were tested so that we are sure to have at least one plate suited for colony counting. Cells from both strains formed a colony on YPD medium, whereas only cells of the *vps5Δ* strain formed a colony on YPD supplemented with hygromycin. We can easily compute the fraction of *vps5Δ* cells from these colony counts. An example of plating is shown in Figure S10.1, corresponding to the strong stringency condition. We observe the clear increase of colonies between 14 and 62 hours on the YPD supplemented with hygromycin.

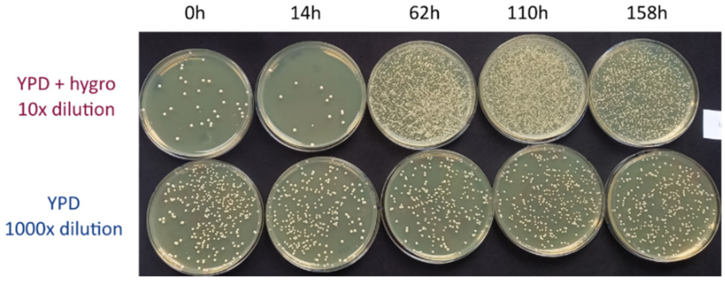

***Figure S10.1****: Culture samples taken from a spike-in experiment under strong stringency, starting with a vps5Δ:parental ratio of 1:1000, are plated onto YPD with or without hygromycin solid medium. We show here the dilutions that are the most illustrative of the enrichment.*

### Digital PCR procedure to compute fraction of *vps5Δ* cells

Digital PCR consists of performing a PCR reaction simultaneously in thousands of microdroplets after random fragmentation of the reaction mix. At the end of the PCR program, each droplet gives a binary signal: amplification or no amplification, depending on the presence of the template in the droplet. Based on the number of droplets with positive amplification, the system computes the number of template molecules per µL of sample. By targeting a template specific to the *vps5Δ* strain and a template common to both strains, we can compute a ratio of templates, which we convert into fraction of *vps5Δ* cells using a calibration curve. Concretely, we targeted the HygR cassette, specific to *vps5Δ* strain, and the *COG6* gene, common to both strains. We grew both strains in monoculture in the bioreactor and took culture samples for genomic extraction. We mixed the two genomes in known ratios and performed digital PCR. It allowed us to obtain the calibration curve giving the relation between the ratio of templates HygR/*COG6* and the known fraction of *vps5Δ* genome (Figure S10.2).

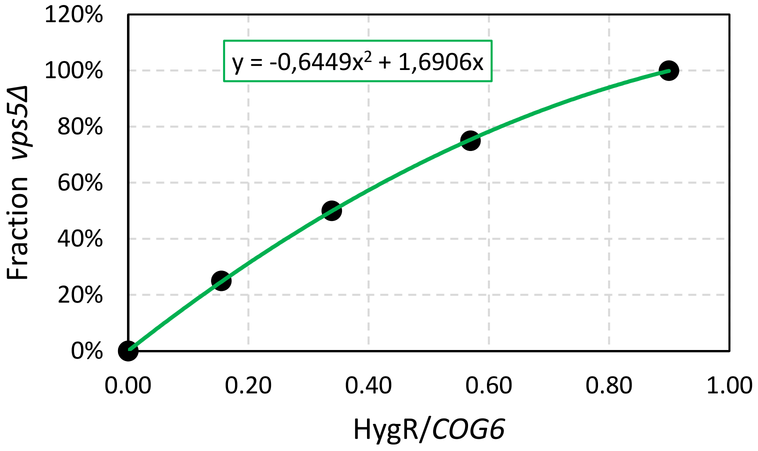

***Figure S10.2**: Samples from the autocrine vps5Δ and parental strains grown individually in the bioreactors are taken for genome extraction. The resulting genomes are mixed in known ratios and used as templates for digital PCRs targeting both COG6 and HygR genes.*

We extracted the genomes from the samples of the spike-in experiment, performed the digital PCRs and obtained HygR/*COG6* ratios. We then used the calibration curve to infer the fraction of *vps5Δ* strains. We included samples from monocultures of both strains and observed that, for samples containing only the parental strain, the assay gave a small percentage of *vps5Δ* (Table S10.1). This does not come from the calibration curve, because positive droplets were also observed in the raw digital PCR data. It could come from unspecific amplification during the PCR and results in limitations regarding the detection threshold. In other words, this renders impossible to interpret the results of co-culture samples with low fraction of *vps5Δ*. We noticed that samples containing only the *vps5Δ* strain gave approximately 100% fraction of *vps5Δ*.

***Table S10.1**: Estimated fractions of vps5Δ cells obtained by dPCR on individual cultures of the parental and vps5Δ strains grown in bioreactor without selection.*

| | Parental strain | *vps5Δ* strain |
|---|---|---|
| **Method** | dPCR | dPCR |
| **0h** | 0.08% | 97.17% |
| **14h** | 0.12% | 98.45% |
| **62h** | 0.10% | 98.56% |
| **110h** | 0.13% | 100.56% |
| **158h** | 0.13% | 96.16% |

We then analyzed samples from the spike-in experiment and compared results obtained with both the plating and the digital PCR procedures (Table S10.2). Both methods gave similar estimations of *vps5Δ* fraction. We computed the relative change between results from the two methods which naturally highlights the differences for the low fractions. Knowing the starting fraction is important to compute the enrichment. We know from Table S10.1 that our digital PCR assay is not suited for estimation of low fractions. Results from the plating procedure are coherent for early timepoints, at 0h and 14h. Indeed, we expect *vps5Δ* fraction to be around 0.1% or lower, because strains were mixed to obtain a *vps5Δ* fraction of 0.1%, and the slightly slower growth of the *vps5Δ* strain could have decreased the fraction between the inoculation and the induction at 0h. Both methods are rather in agreement for higher fractions, confirming the enrichment of the *vps5Δ* strain.

***Table S10.2**: Fraction of vps5Δ cells obtained by Plating and dPCR for co-cultures started with a vps5Δ:parental ratio of 1:1000, grown in bioreactors under mild or strong stringency.*

| | Mild stringency | | | | | |
|---|---|---|---|---|---|---|
| | rep1 - reactor 9 | | | rep2 - reactor 11 | | |
| **Method** | Plating | dPCR | relative change | Plating | dPCR | relative change |
| **0h** | 0,07% | 0.20% | 186% | 0.07% | 0.48% | 586% |
| **14h** | 0.08% | 0.19% | 138% | 0.06% | 0.23% | 283% |
| **62h** | 1.67% | 2.19% | 31% | 1.71% | 2.31% | 35% |
| **110h** | 30.60% | 32.56% | 6% | 30.36% | 34.36% | 13% |
| **158h** | 19.33% | 20.41% | 6% | 12.15% | 14.06% | 16% |

| | Strong stringency | | | | | |
|---|---|---|---|---|---|---|
| | rep1 - reactor 13 | | | rep2 - reactor 15 | | |
| **Method** | Plating | dPCR | relative change | Plating | dPCR | relative change |
| **0h** | 0.09% | 0.24% | 167% | 0.07% | 0.16% | 129% |
| **14h** | 0.05% | 0.16% | 220% | 0.05% | 0.17% | 240% |
| **62h** | 42.74% | 45.10% | 6% | 48.10% | 49.90% | 4% |
| **110h** | 29.73% | n.a. | n.a. | 22.99% | 26.24% | 14% |
| **158h** | 6.59% | 8.10% | 23% | 2.04% | 2.63% | 29% |

**Capacity of FACS-based screening to enrich the *vps5Δ* strain**

We wanted to test whether cells with higher downstream activation could be enriched through FACS sorting. To investigate this question, a spike-in experiment was conducted with the parental and *vps5Δ* autocrine strains, starting with a *vps5Δ* fraction of 0.1%. The co-culture was grown in batch culture with the OptoTube setup with 100% light DC. At the end of the culture, cells were harvested and resuspended in cold PBS for FACS sorting based on downstream activation (mScarletI fluorescence). The gating strategy is presented in Figure S10.3, in which the P4 population is sorted.  We recovered the sorted P4 population and the pre-sorting population, which represents the population at the end of the culture, before the sorting. They were then grown overnight in liquid YPD and plated on solid YPD medium, supplemented or not with hygromycin. Colonies were counted to establish the fraction of *vps5Δ* cells. Sorting allowed approximately 40-fold enrichment in the sorted population compared to the pre-sorting population (Table S10.3). We conclude that cells with higher downstream activation can be enriched through FACS, but that growth-based selection is much more efficient than one round of FACS.

***Table S10.3****: Fraction of vps5Δ cells in pre-sorting and FACS-sorted population after a batch co-culture started with 1:1000 vps5Δ cells.*

| | **Fraction of *vps5Δ* Pre-sorting population** | **Fraction of *vps5Δ* Sorted population** | **Enrichment** |
|---|---|---|---|
| Replica 1 | 0.057% | 2.076% | **36.4** |
| Replica 2 | 0.043% | 2.033% | **46.9** |

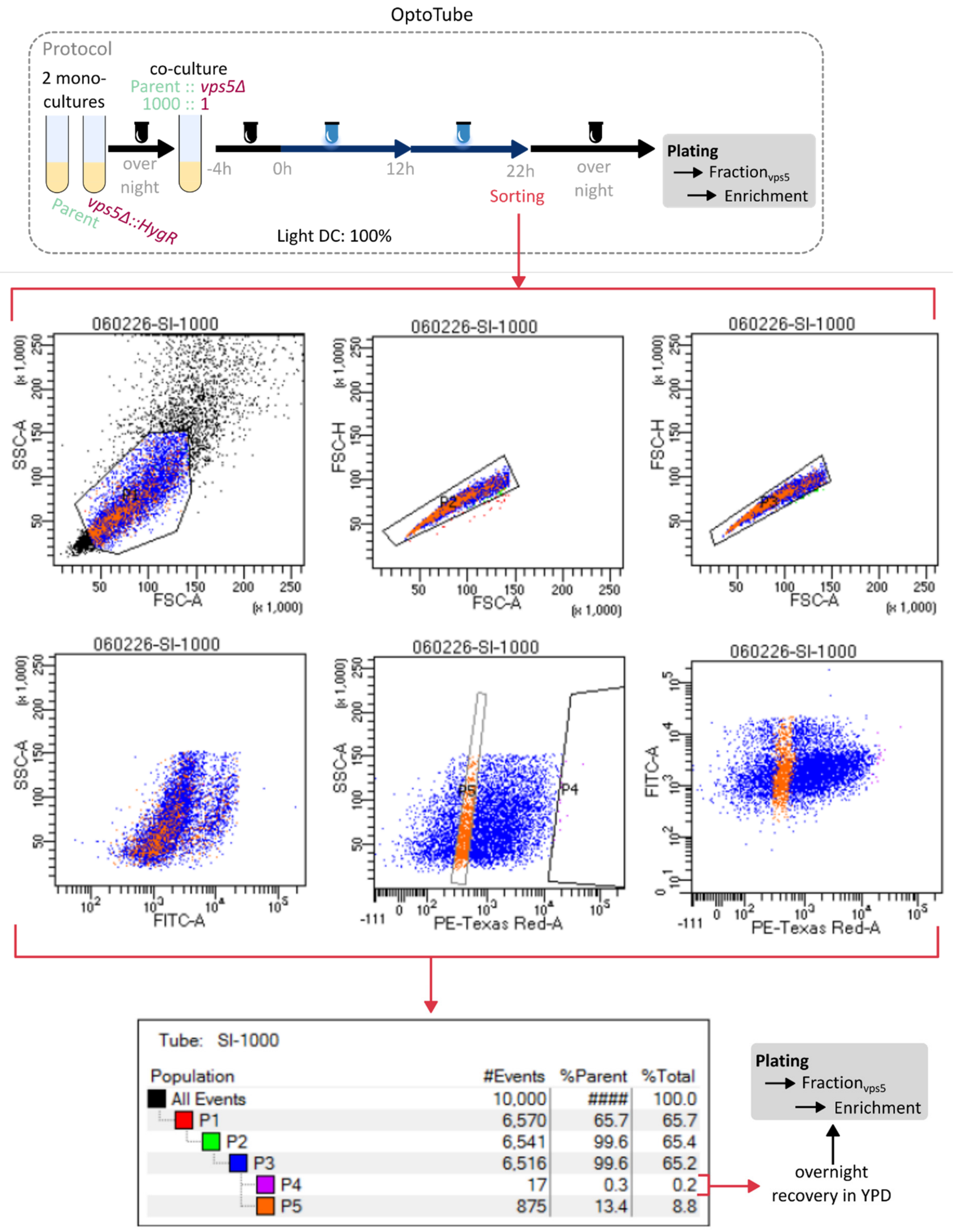

| Population | #Events | %Parent | %Total |
|---|---|---|---|
| All Events | 10,000 | #### | 100.0 |
| P1 | 6,570 | 65.7 | 65.7 |
| P2 | 6,541 | 99.6 | 65.4 |
| P3 | 6,516 | 99.6 | 65.2 |
| P4 | 17 | 0.3 | 0.2 |
| P5 | 875 | 13.4 | 8.8 |

***Figure S10.3**: Assessment of FACS-based screening with the AutoGrowth biosensor. The vps5Δ and parental strains are mixed in ratio 1:$10^3$ and grown in the OptoTube setup in his+ medium under 100% blue light duty cycles. At the end of the second culture phase, at 22h, co-cultures are sorted following a gating strategy to obtain the sorted population P4 with cells showing the highest signal in the red channel. Sorted and pre-sorting populations are then grown overnight and plated for colony counting.*

### Supplementary Note 11: Investigation of the cheater behavior

During the spike-in experiment presented in Figure 5B, the decrease of the *vps5Δ* fraction observed under strong stringency at days 4 and 6 was not expected. We made the hypothesis that it could come from parental cells gaining the ability to grow despite the selection pressure, that is, a cheater behavior. To investigate this question, we leveraged culture samples plated on YPD and YPD+hygromycin medium from the spike-in experiments under strong stringency (Figure 5B top, Supplementary Note 10). We used clones coming from the 158h timepoint, that is, after 6 days of culture. Six isolated colonies (Y1-6) were taken from YPD medium, and three from YPD supplemented with hygromycin (H1-3). None of the Y1-6 clones grew on YPD hygromycin, indicating that they were parental cells. These clones were tested under 100% light duty cycles in the OptoTube setup (Figure S11.1A and B). First, as expected, secretion levels of Y1-6 and H1-3 clones corresponded to those of the parental and *vps5Δ* strains respectively. It indicates that no adaptation comes from changes in the secretion capacity. More surprisingly, downstream activation levels were similar for the two strain types, and, in both cases, higher than the parental and *vps5Δ* strains. Although there is variability among the Y1-6 clones, the average activation level represents an 8-fold increase compared to the parental strain (see Figure S8.1). The increase is smaller for the *vps5Δ* strain, with 1.3-fold only. It confirms that cells with upregulated downstream activation arise, which allows them to cope with the stringency despite unchanged secretion levels. Interestingly, several Y1-6 clones even showed downstream activation in the absence of light, suggesting that the reporter tends to be constitutively expressed in cheater cells (Figure S11.1C).

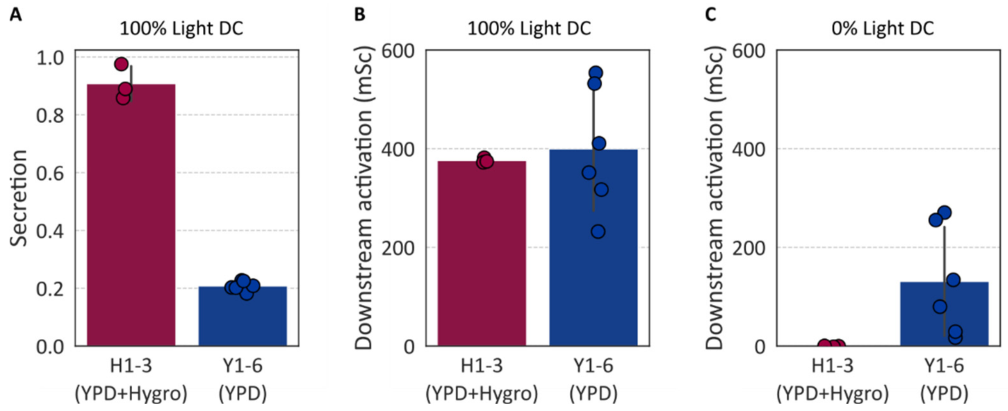

***Figure S11.1***: *Cheater cells up-regulate downstream activation.* ***A) and B)*** *Y1-6 and H1-3 clones are grown in OptoTube setup in his+ medium for 4 hours in the dark then for 12 hours under 100% blue light DC, before inoculation of fresh his+ medium for 10 hours under blue light, followed by flow cytometry for measurements of cell fluorescence and secretion.* ***C)*** *Y1-6 and H1-3 clones are grown in OptoTube setup in his+ medium for 16 hours in the dark, before inoculation of fresh his+ medium for 10 hours in the dark, followed by flow cytometry for measurements of cell fluorescence. Experiments are done with three or six biological replicates; medians of each population are represented by dots, and their mean and the corresponding standard variation is shown with the bar plot*

The fact that the *vps5Δ* clones also showed upregulated activation indicates that cells from both strains adopted a "cheater" behavior. In addition, we observed in Figure 5B top that downstream activation stabilizes at two different levels for mild and strong stringencies. This would suggest that medium stringency defines a selection pressure, which represents for cells a level of activation to reach to be able to cope with this pressure. This leads to upregulation of downstream activation, which uncouples growth rate and secretion capacity. Consequently, strain fitness becomes the only driver of selection, resulting in a decrease of *vps5Δ* fraction due to its altered fitness compared to parental strain (observed in Figure 5B). We also note that cheater cells only have upregulated activation to cope with selection, but do not cheat by lower scFv secretion.

An additional control experiment was performed by growing parental strain alone under strong stringency for several days. We observed a growth decay after the switch to selective medium at 0h (Figure S11.2). Importantly, the population started recovering after 2 days and reached normal growth after 4 days. It means that the appearance of cheater cells does not depend on a cross-feeding mechanism between strains, but that it is inherent to the selection system. Hence, screening under strong stringency should not last more than 2 days.

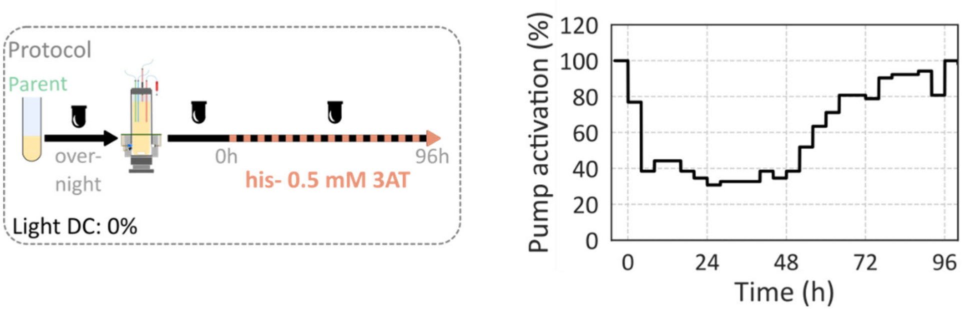

***Figure S11.2**: Cheater behavior is independent of cross-feeding mechanisms. The parental strain was grown in eVOLVER bioreactor in turbidostat in the dark at first in 40% histidine medium, before switching at time 0h to his- medium supplemented with 0.5 mM 3AT.*

### Supplementary Note 12: Automated analysis pipeline of flow cytometry data

#### Processing flow cytometry data of cell fluorescence

Our automated analysis pipeline to go from raw fluorescence data to fluorescent protein levels (in arbitrary units) is presented in Supplementary Figure S12.1A. It is based on a previously published pipeline[28]. The entire pipeline was performed using Python scripts. For gating, a first threshold was set to keep only cells with the forward scatter height (FSC-H) values above 250. Then, we kept cells in the dense region of forward and side scatter data (Figure S12.1B), and removed doublets based on deviation from linearity between the Area and Height of the forward scatter (Figure S12.1C). This procedure selected 68%-78% of the events. For co-cultures, gated data were then subjected to automated clustering based on log(BLU-V/FSC-H) to classify each cell as an autocrine or paracrine strain (Figure 2A). Next, spectral deconvolution was performed by first removing autofluorescence and then computing the contribution of each fluorescent protein to the resulting signal of the cell (Figure S12.1D). This estimated contribution represents the amount of fluorescent proteins expressed by the cell in arbitrary units.

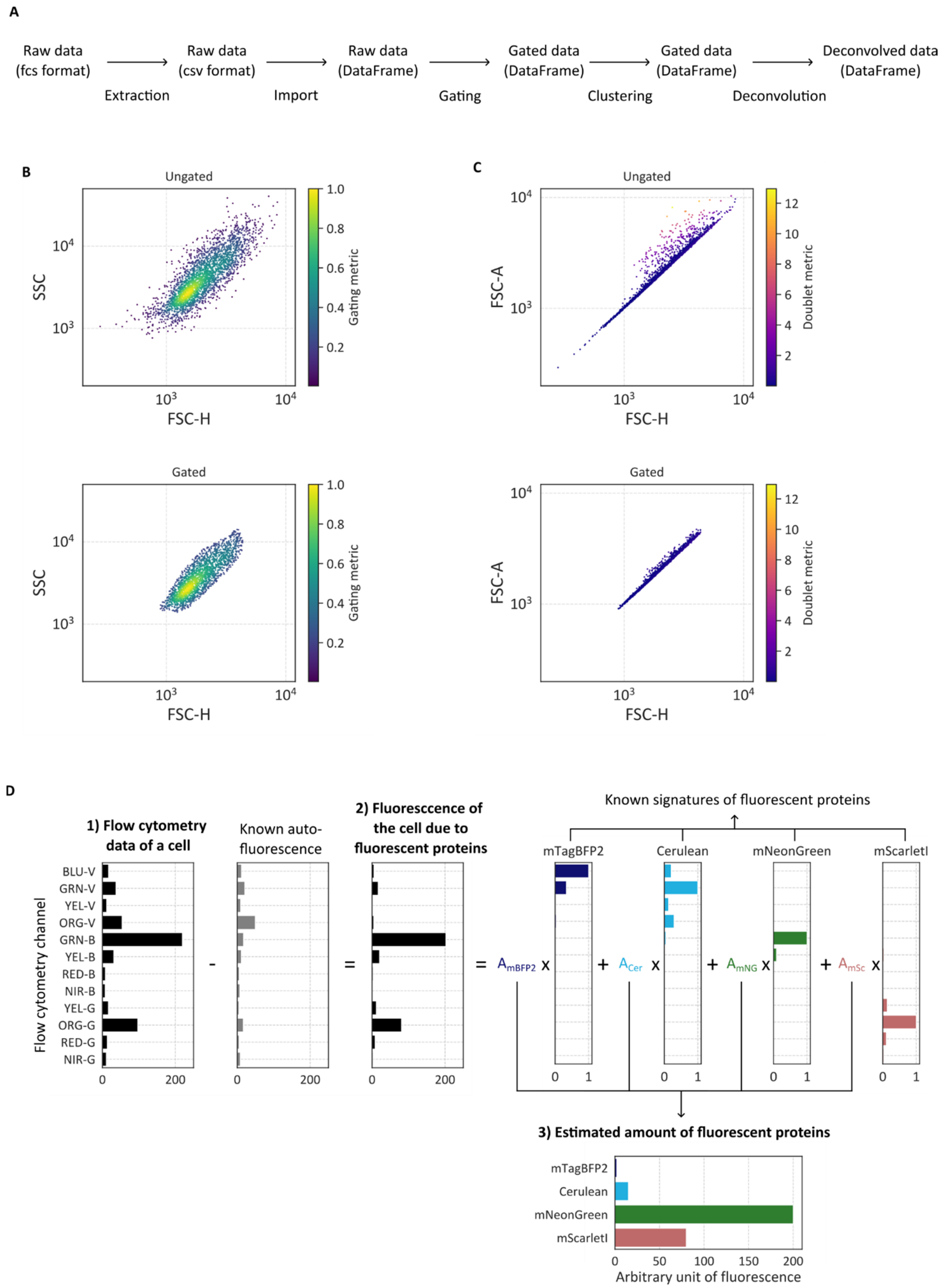

***Figure S12.1**: Automated pipeline for processing flow cytometry data of yeast cell fluorescence.* ***A)*** *Flow of the pipeline.* ***B)*** *Gating based on forward and gate scatter to keep only dense region (gating metric < 0.2).* ***C)*** *Removal of doublets using ratio between area and height of forward scatter (doublet metric <1).* ***D)*** *Deconvolution strategy to infer amount of fluorescent protein in a cell, in arbitrary fluorescence units.*

**Processing flow cytometry data of bead fluorescence**

In this study, the protein secretion measurements were performed using anti-Flag magnetic beads that capture the Flag-tagged protein of interest, whose amount can then be quantified thanks to its fusion with mNeonGreen. We used the approach documented in Sosa *et al*, 2023[8]. The pipeline used to process beads data is presented in Figure S12.2A. As for the cells, tasks are all performed in Python. After extraction and import, gating procedure is applied to keep only beads of sufficient size (green population, Figure S12.2B). More specifically, a first threshold is applied to keep only cytometer events with a forward scatter higher than 500, and a second threshold based on both forward and side scatter is applied to remove cells and keep only beads. Next, data are processed to obtain the secretion readouts. As internal controls, Flag-tagged Cerulean proteins are added during the incubation of the beads with the sample to account for the variations of bead size and binding capacity. We ran in parallel a tube containing only Cerulean, called Cerulean sample, used for normalization. We illustrate the pipeline with data from Figure 2B on secreted mNeonGreen, in which the autocrine strain secreting mNeonGreen is induced with various light duty cycles. First, we normalized the fluorescence obtained from mNeonGreen (GRN-B-HLin, Figure 12.2C, top left) by the fluorescence obtained with Cerulean (BLU-V-HLin, Figure 12.2C, top right) for each bead. It gave us the ratiometric signal GRN-B/BLU-V (Figure 12.2C, bottom left). Finally, the median ratiometric signal of the Cerulean sample was subtracted from that of each sample to isolate the signal originating solely from the bead-captured protein of interest (Figure S12.2C, bottom right).

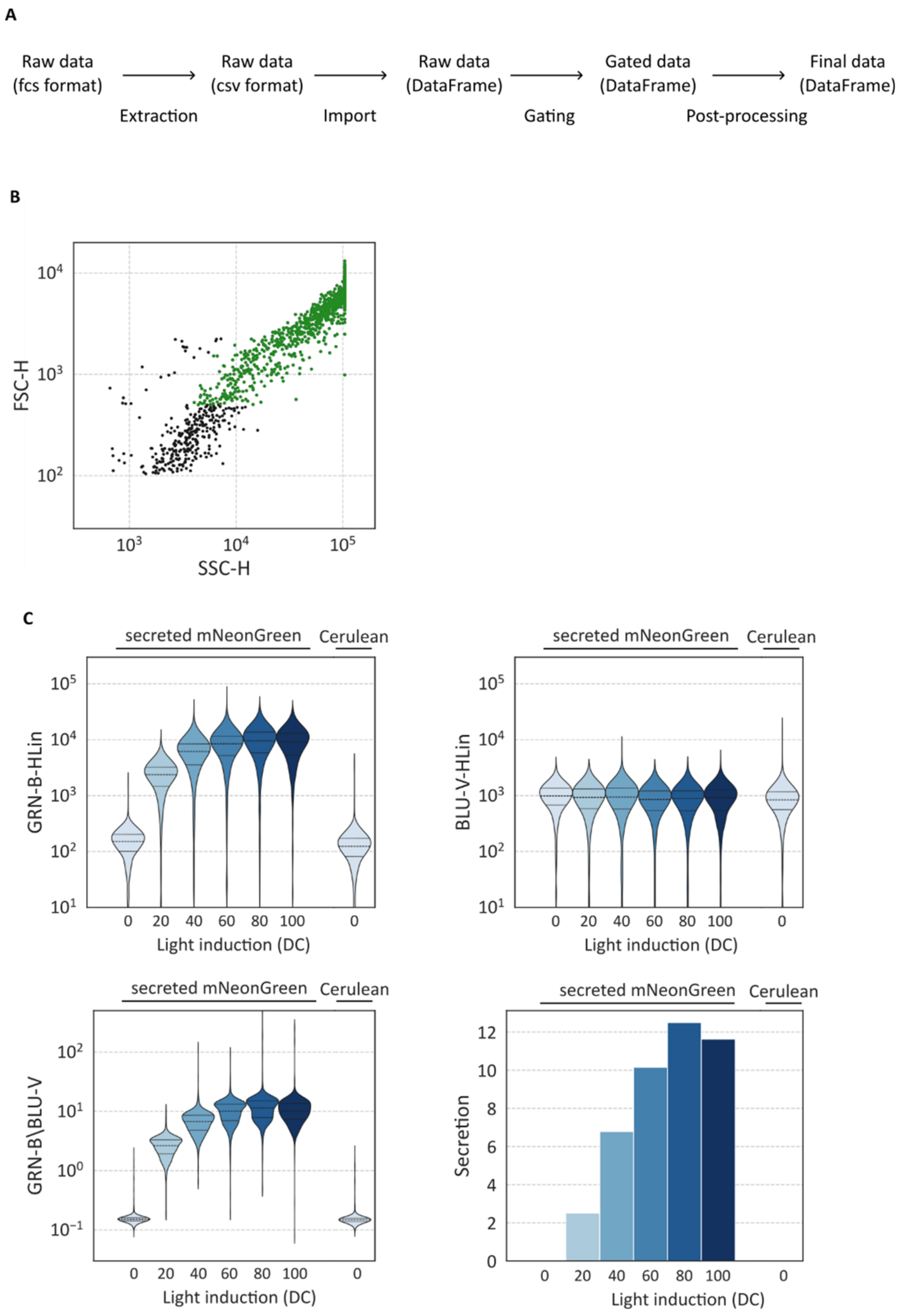

***Figure S12.2**: Pipeline for processing flow cytometry data of bead fluorescence. **A)** Flow of the pipeline. **B)** Gating based on forward and side scatter to keep only beads (green dots). **C)** Post-processing of the bead fluorescence, with the use of GRN-B-HLin and BLU-V-HLin data to compute the ratiometric GRN-B/BLU-V.*
